# Variability and scale-dependence in molecular evolution rate impacts interpretation of eukaryotic evolutionary histories

**DOI:** 10.64898/2026.05.25.724440

**Authors:** Erik Tamre, Lyle Nelson

## Abstract

Molecular evolution is often modeled as proceeding at consistent rates over time, with some deviations accommodated by relaxed molecular clock models. Here, we quantify the full extent of variability in branch-specific substitution rates across relatively well-calibrated eukaryotic phylogenies, confirming that punctuated change at the molecular level underlies evolution at the morphological level where punctuated dynamics are more commonly recognized. We also show how inferred substitution rates decrease systematically when measured across increasing time intervals. This scale-dependence persists across alternative clock models, calibration strategies, and prior assumptions, but disappears in simulated data evolved under a constant rate - suggesting that the phenomenon arises from time-varying substitution rates and reflects genuine properties of evolutionary histories rather than model artifacts. The observed pattern is analogous to the Sadler effect in sedimentary geology, where time-averaged rates decline with increasing measurement interval because sedimentation is episodic, with longer hiatuses occurring less frequently. The recognized scale-dependent bias in molecular evolution is not captured in current molecular clock models and significantly impacts inferences of evolutionary history, such as estimating the age of Metazoa and understanding the timing and nature of the Cambrian Explosion.

**Significance Statement:** Rates of molecular evolution are central to reconstructing the history of life, yet they are often assumed to be approximately constant over sufficiently long timescales. By analyzing relatively well-dated evolutionary trees of eukaryotes, we show how inferred rates of genome change systematically decrease as the timescale of measurement increases. This pattern is analogous to a well-known phenomenon in sedimentary geology where apparent sedimentation rates decline over longer intervals due to episodic processes. Our results demonstrate how long-term variability in evolutionary rates similarly produces a significant scale-dependent bias which is overlooked in current evolutionary models. Recognizing and quantifying this effect is important for dating key evolutionary events, such as the origin of animals, and for understanding the cadence of evolutionary processes.

## Introduction

The paleontological record shows that evolution at the morphological level varies in rate over time, inspiring views such as punctuated equilibria (1, 2) where periods of relative evolutionary stasis are interrupted by shorter intervals of rapid change. However, the question of the changing tempo and mode of evolution has received much less explicit consideration at the molecular level. If anything, a more regular process is often assumed there – such as in molecular clock approaches, which take pains to model rate variability but also rely on at least some tendency towards clock-like behavior in sequence evolution. The accuracy of this view over geological time is challenging to test because no sequence data survives from the geologically distant past: useful sequence is not usually recovered from DNA and proteins older than ∼1 Ma, with only a few exceptions of extraordinary preservation extending back to the Miocene (3). Molecular phylogenies allow estimates of the amount of sequence evolution between the nodes of a phylogeny, but time-calibrating the phylogeny – usually based on dated taxonomically diagnostic fossils from the rock record – is a challenging problem in most clades given the scarcity of such a record. Since the proposal of a molecular clock by Zuckerkandl and Pauling (4), the relationship between absolute time and evolutionary change has thus commonly been approached from the other direction (5): the timing of the nodes in a phylogeny is estimated by assuming a somewhat predictable, if variable, rate of evolution, which then allows the translation of evolutionary distances into absolute time.

However, paleontological advances and especially the increasing availability and precision of radiometric ages that date fossil occurrences now make it possible to measure substitution rates in the geological past for at least some clades with documented, well-dated fossil records – largely eukaryotes, and particularly macroscopic organisms with a high preservation potential such as animals and land plants. Studying substitution rate variability is important for our understanding of evolutionary processes, but also for assessing whether they are captured by practical phylogenomic tools such as molecular clock models. Some previous molecular clock studies have assessed the molecular clock hypothesis (predictable rates of substitution) by plotting the amount of cumulative sequence divergence against node age as constrained by the fossil record; this practice was especially common early in the history of the field, so as to demonstrate the applicability of the technique for constraining divergence times in the specific dataset (e.g., (6)). This test can show that the cumulative increase in sequence change tends towards a mean rate, but lacks sensitivity to variations in rate on individual branches – especially ones traversing only short amounts of time and therefore having limited impact on cumulative divergence regardless of rate. Lee et al. (7) calculated branch-specific rates of morphological as well as molecular evolution in arthropods and identified elevated rates around the Cambrian Explosion, but Paterson et al. (8) found constant branch-specific rates of morphological evolution across the entire Cambrian in a record of trilobites. Bokma (9) explicitly suggested the use of molecular phylogenies to look for punctuated equilibria, but examined changes in speciation and extinction rates and cladogenesis rather than measuring substitution rates. Pagel et al. (10) observed gradual and punctuational components in a record of molecular evolution, but did not attempt to measure rates by calibration to absolute time. Recognizing the presence of these components in the evolutionary process, Douglas et al. (11) have recently developed a sophisticated probabilistic framework allowing punctuated as well as gradual change to be modelled in the analysis of genomic, morphological, as well as linguistic data.

Certain kinds of discontinuous rates can also give rise to scale-dependent observation biases, such as long recognized in sedimentary geology as the Sadler effect (12). The Sadler effect describes the phenomenon whereby stratigraphic packages yield progressively lower time-averaged sedimentation rates when measured over longer intervals. This phenomenon arises because sedimentation is episodic and contains hiatuses (with longer hiatuses less frequent), creating a fractal scaling relationship where the time span of the measurement and the observed sediment accumulation rate can be related by an empirical negative power law (12, 13). Gingerich (14) suggested that this effect may also extend to rates of morphological evolution. More recently, such scale-dependence has been reported in a variety of evolutionary rates (15, 16) – with the evidence in molecular evolution (e.g., (17)) particularly focused on the mutation-substitution transition, which describes the contrast between high mutation rates observed in short-term evolution studies and much lower rates of gene fixation at the population scale and above.

Thus, in sedimentary geology, rates are considered in the context of measurement timescale, and there is evidence suggesting that need also for rates of evolution – but substitution rates are not considered in this context in molecular clock studies. In this study, we quantify the rate variability of molecular evolution in eukaryotes and demonstrate the size and impact of the resulting Sadler-like effect over timescales of up to a billion years, well beyond the mutation-substitution threshold. We explicitly measure the variability of molecular substitution rates on individual branches (covering millions of years) in the eukaryote phylogeny across multiple genomic datasets and fossil calibration schemes that have been proposed for constraining their evolutionary history. We report considerable (over an order of magnitude) substitution rate variability even in genes chosen for their clock-like behavior and within eukaryotic clades with well-dated fossil records. We quantify the resulting scale-dependent observation bias and demonstrate its implications for both understanding the evolutionary process and constraining divergence times within specific clades such as Metazoa.

## Results

### Exploring substitution rate variability

We focused on estimating substitution rate variability in a eukaryotic dataset previously assembled for a molecular clock study (18), taking advantage of the relatively high number of time-constrained branches afforded by the broad sampling and the dated fossil record of multiple clades of eukaryotes. The inferred substitution rates on each branch (see **Methods**) are shown in **Figure 1a**.

**Figure 1.**
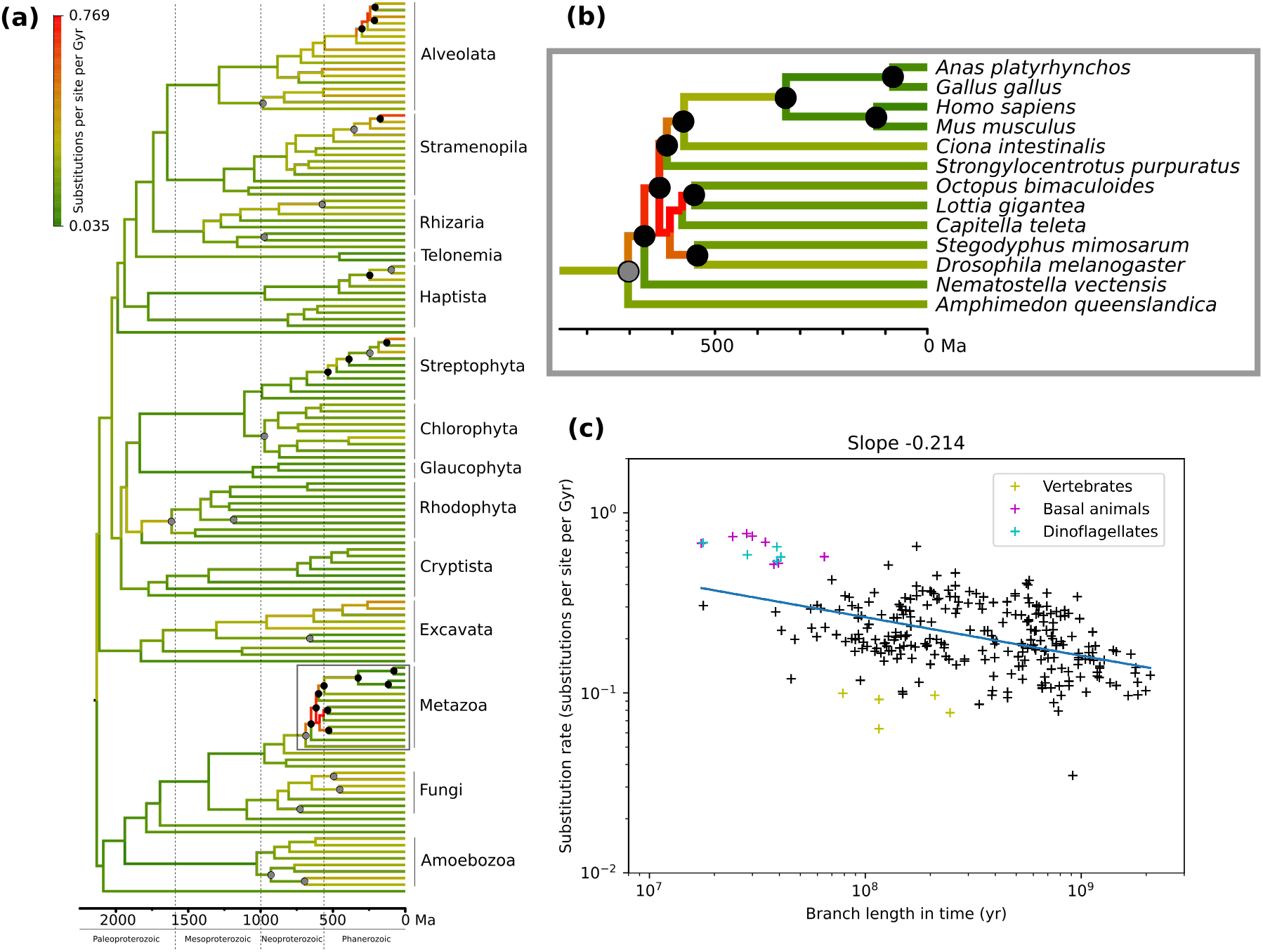
**a)** Rate variability of molecular evolution among eukaryotes inferred from a molecular clock in (18). Branches are colored according to the average substitution rate inferred along that branch using an autocorrelated relaxed clock model, with the tree rooted on Amorphea. Black circles indicate nodes with tight time-calibration (bracketed within 150 Myr or less) from the fossil record, and grey circles indicate nodes calibrated with broader (>150 Myr) constraints. The chronogram with full taxon labels is shown on **Figure S1**. **b)** Corresponding scatter plot showing the relationship of branch length (between mean posterior node ages) and the inferred substitution rate on each branch. Least-squares fit is shown in blue and its slope displayed at the top. The negative slope reflects a Sadler-like effect where lower rates are observed on longer branches. High rates among early animals are shown in magenta and among dinoflagellates in cyan, with low rates among vertebrates in yellow. Note that while these are the best-constrained clades in terms of fossil calibrations, they do not drive the whole effect, since their exclusion still results in a slope of -0.16. **c)** Metazoan part of the tree magnified from panel **a**. Note that both the fastest and slowest substitution rates inferred on the eukaryote tree are found within Metazoa, likely due to the highest density of fossil calibrations there.

Clusters of highest and lowest substitution rates are inferred in parts of the tree where branches are bracketed by calibrations: for example, close to base of Metazoa (high rates), within dinoflagellates (high rates), and within vertebrates (low rates). The mean of all branch rates on the tree is 0.237 substitutions per site per Ga. Internal branches close to the base of crown Metazoa after the divergence of sponges display rates of 0.52-0.77 substitutions per site per Ga (with the upper end being the highest inferred on the whole tree), while internal branches within dinoflagellates after the divergence of Noctilucales show inferred rates of 0.54-0.68 substitutions per site per Ga. Conversely, branches within crown vertebrates – represented here by birds and mammals – show rates of 0.06-0.10 substitutions per site per Ga, in agreement with previous observations of low substitution rates in vertebrates (6). This represents an order of magnitude of variability in inferred substitution rates averaged across a large set of genes chosen for clock-like behaviour, with both extremes realised in Metazoa (see **Figure 1b**) – a relatively small but well-calibrated clade within eukaryotes. We also found a similar degree of rate variability among animal lineages in an alternative dataset from (19), with highest rates again appearing close to the base of Metazoa (see **Supporting Text**).

To understand whether the observed substitution rate variability is driven by specific genes or reflects genome-wide effects, we further examined rate variability in the dataset from (18) on a gene-by-gene basis – focusing on Metazoa as the best-calibrated part of the tree, and also where the rate averaged across all genes varied most significantly. Across branches in Metazoa, most genes in the dataset show a standard deviation of branch rates amounting to 1-2 times the mean rate for each gene (**Figure 2a**), reflecting that each gene’s history included significant and comparable amount of rate variability within this subtree. Furthermore, rate changes in different genes show a degree of correlation to each other: pairwise regression between rates on corresponding branches in two different genes (see **Figure S2** for an example) yields a positive R-value for over 85% of gene pairs in this dataset, with the mean R-value being 0.31 (histogram on **Figure 2b**).

**Figure 2.**
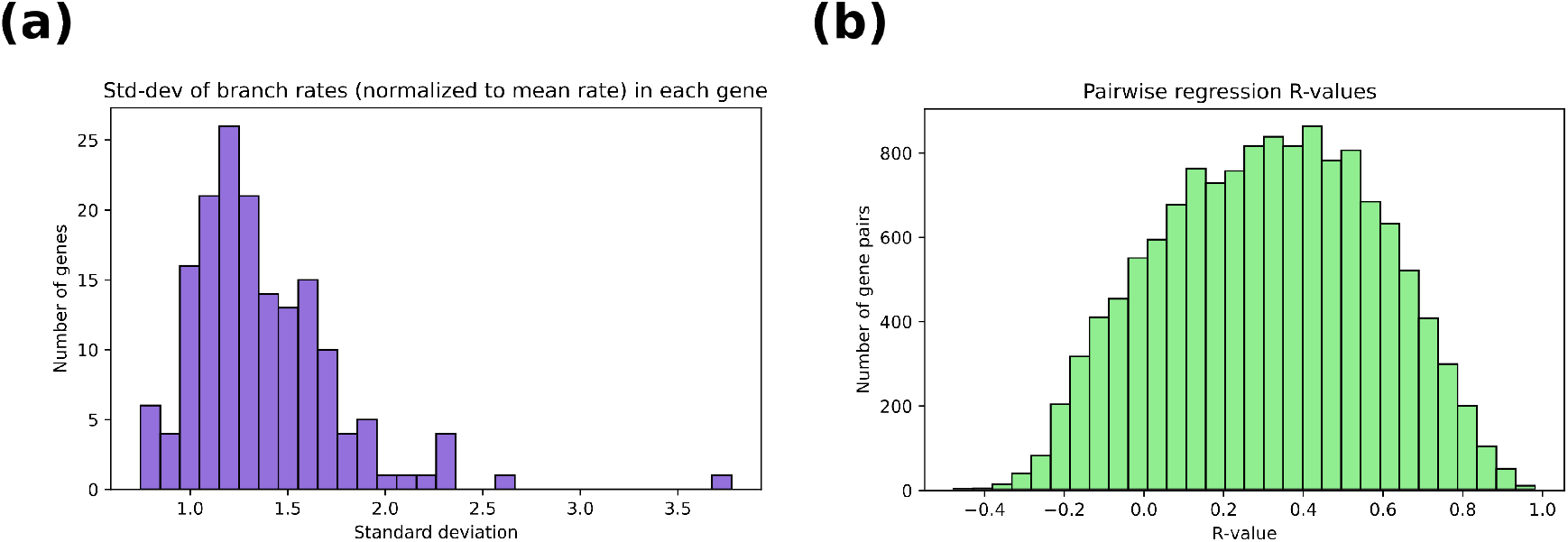
**a)** Histogram showing the gene-by-gene standard deviation of substitution rates across the metazoan part of the tree, normalized to the mean rate within each gene. Note that there are no particularly clock-like genes – for every gene, the standard deviation of branch rates is close to the mean rate or higher. **b)** Histogram showing R-values for pairwise regressions between rates on corresponding branches in the metazoan part of the tree for each pair of included genes. Note that the distribution is centered well above zero, suggesting some correlation in time-dependent rate changes across the genome.

### Scale-dependence in substitution rate estimates

To test for a Sadler-like scale-dependence of inferred substitution rates stemming from the observed rate variability in the dataset from (18), we plotted on a log-log scale the substitution rate on each branch against the length of the branch (**Figure 1c**, cf. Figure 1 in (12)). There is a clear negative relationship between these two variables, with the slope of the regression line at -0.214 (p-value ≈ 10^-15^ for a test whether the slope is zero). Given the log-log scale, the units of the slope are “orders of magnitude of change in inferred substitution rates per order of magnitude of change in the measurement timescale”. In other words, a slope of -0.214 corresponds to a 2.7x reduction in average substitution rates upon an increase in measurement timescale by two orders of magnitude from 10^7^ years to 10^9^ years. We found an even stronger scale-dependence in the alternative dataset focused specifically on Metazoa (19), yielding a slope of -0.308 or -0.295 depending on tree prior choice (birth-death or uniform, respectively) – see **Supporting Text**.

These slope values understate the true size of the scale-dependent observation bias, since some branches on a molecular clock, far away from fossil or present-day calibrations, are driven almost entirely by the clock model – which penalizes departures from the mean rate and does not allow for independent rate measurements. To mitigate that issue, to estimate the true magnitude of the scale-dependence, and to better understand its sources, we re-ran the molecular clock analysis on the dataset from (18) with variations in molecular clock setup, including without sequence data.

### Testing the impact of sequence data, internal node calibrations, and tree prior

We re-analyzed the dataset (see **Methods**) by making relaxed molecular clocks under different combinations of 1) including or omitting sequence data (running the clock under the prior only), 2) including or omitting fossil calibrations on internal nodes, and 3) tree prior choice (birth-death or uniform) – see **Table 1**.

**Table 1.**
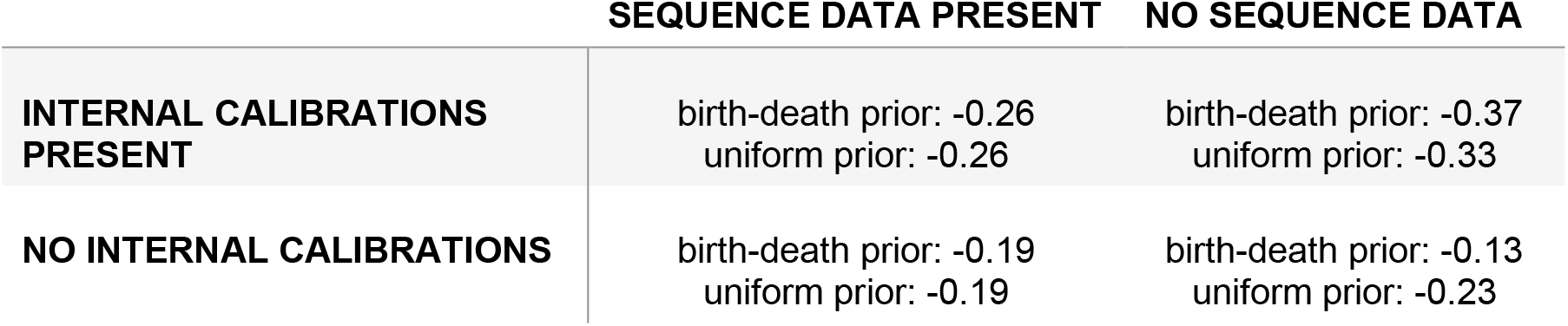
Strength of the observed Sadler-like effect under the presence/absence of sequence data, presence/absence of internal calibrations, and choice of tree process prior for the dataset from (18). The strength of the Sadler-like effect is quantified by slope value from a linear regression on a log-log scale, showing orders of magnitude of change in inferred substitution rate per order of magnitude of change in branch length.

The removal of sequence data under the birth-death prior results in the Sadler-type plot with a slope of -0.37 (**Figure 3b**), corresponding to a 5.5-fold reduction in average substitution rates upon an increase in the measurement timescale by two orders of magnitude. Since prior-only results should be more sensitive to tree prior choice as there is no sequence data to overwhelm the prior, we also tested a uniform tree prior which returned a slope of -0.33 (**Figure S3**), corresponding to a 4.6-fold reduction in average substitution rate over two orders of magnitude in branch length. Predictably, prior choice did not play as significant a role in re-runs including sequence data: birth-death (**Figure 3a**) and uniform (**Figure S4**) tree priors both yielded a slope of -0.26, with the clock model driving rates towards the mean and flattening the slope compared to prior-only runs (**Figure 3b, Figure S3**).

**Figure 3.**
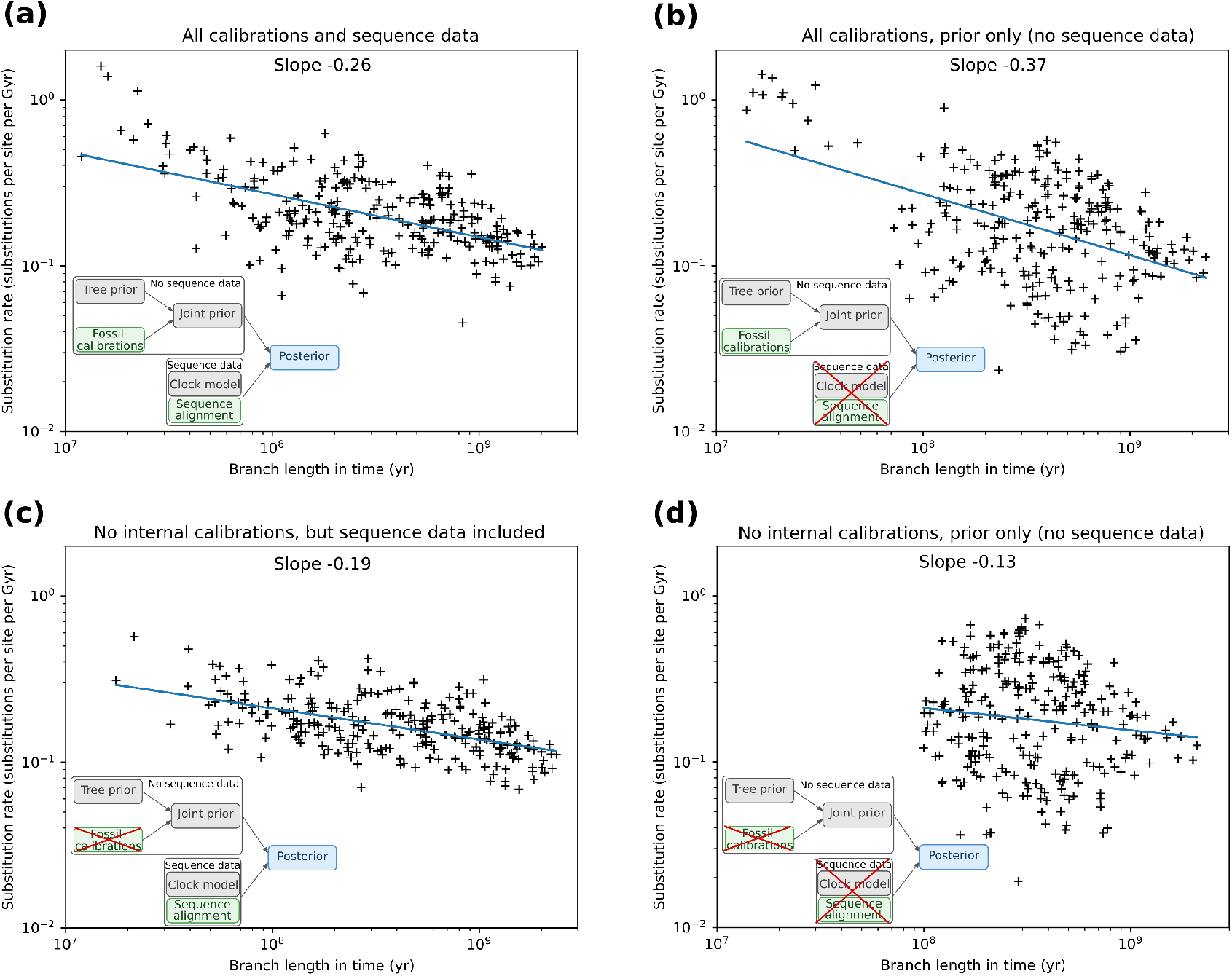
The Sadler-like scale dependence of substitution rates in the dataset under variations in molecular clock setup. Note that while the fossil calibration data is included at the prior stage, sequence data is only incorporated upon the transition from prior to posterior (see **Figure S7** for a simplified schematic of the Bayesian molecular clock workflow, here depicted on insets). **a)** shows the dependence of inferred substitution rate on branch length in a clock including both internal calibrations as well as sequence data **b)** includes all calibrations but no sequence data, so the posterior is identical to the joint prior and the Sadler-like effect is not understated by the clock model; **c)** includes the sequence data but no internal calibrations, and **d)** omits both internal calibrations and sequence data. The corresponding chronograms are shown in full on **Figures S8-S11**. All panels show runs under the birth-death prior; **Figures S3-S6** show plots for runs under the uniform prior.

A birth-death tree prior (yielding the more pronounced scale-dependence) is likely more appropriate for this tree whose nodes represent speciation events, and the uninformative uniform prior is often dismissed since no obvious evolutionary process would result in branching events thus distributed (20). Regardless, with either prior, the runs without a clock model suggest an approximately five-fold reduction in average substitution rates upon an increase in measurement timescale by two orders of magnitude. The prior-only runs constitute our best effort to measure rates without simultaneously driving them towards the mean using the clock model, and the resulting estimate may be the most appropriate reflection of the true size of the Sadler-like effect in the evolutionary history of eukaryotes.

We also tested the effect of removing all internal calibrations. With sequence data included, the clock model drives rates towards the mean as predicted in the absence of constraints from internal calibrations, resulting in a flattening of the slope to -0.19 with both birth-death and uniform priors (**Figure 3c** and **Figure S5**, respectively). The runs omitting both internal calibrations and sequence data (birth-death: slope -0.13, **Figure 3d**; uniform: slope -0.23, **Figure S6**) have lost most information about the actual evolutionary history, simply displaying the dynamics of the prior (which is why prior choice makes such an impact in these runs, as priors distribute nodes in time differently) – but they nevertheless reflect that timing information from internal calibrations greatly adds to the Sadler-like effect.

In all cases, note that a negative slope is preserved and rates are not fully driven to the mean – even where the sequence data is included and the clock model acts without constraints from internal calibrations. This is likely due to *terminal* calibrations (all sequences are sampled in the present), which constitute another source of real timing information that it does not make sense to remove. While they only constrain the tree on one end, they are still abundant – even this relatively well-calibrated tree has only 33 internal calibrations but 136 terminal calibrations – and fix nodes in time accurately and precisely, rather than to a broad timespan. Terminal calibrations can also provide timing information about branches that are far from internal calibrations, making it worthwhile to include even such branches in the analysis of rate variability.

### Simulated alignment evolved under a constant rate

While all the above analyses retain enough real evolutionary timing information to explain their recovery of the Sadler-like scale-dependence, it is further useful to test that these analysis techniques would recover no such effect in a genomic dataset without any genuine scale-dependence in the evolutionary history of the sequences. To that end, we evolved a synthetic sequence alignment (see **Methods**) across the same underlying eukaryotic timetree at a constant substitution rate, thus expected to create no Sadler-like effect. We then ran our molecular clock analysis on that dataset – without calibrations, as this represents a fictional evolutionary history.

The molecular clock analysis correctly infers that there is little rate variability among the branches, and indeed the inferred Sadler-like effect on substitution rates vanishes (**Figure 4**). This further confirms that the effect is a feature arising from the actual evolutionary history, rather than an artifact of the rate inference technique: e.g., baked into the priors, or resulting from mistakes in implementing the molecular clock. The rate variability and Sadler-like effect are identified in the real molecular data, but not in the control dataset (with no rate variability) that was analyzed using the same technique.

**Figure 4.**
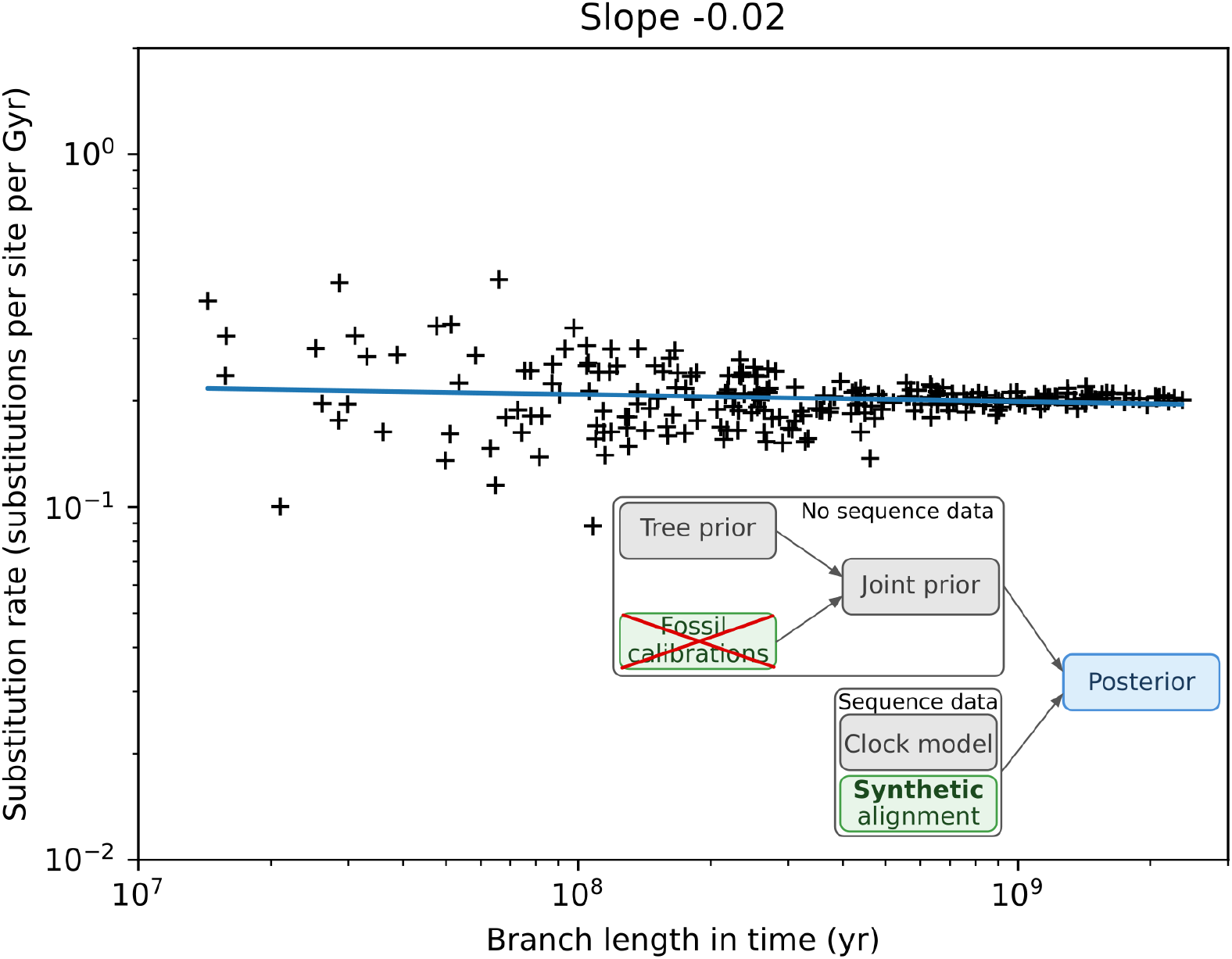
Inferred substitution rate vs. branch length in a synthetic sequence dataset evolved at a constant substitution rate along branches of the molecular clock on Figure 1. The analysis correctly infers the fixed rate, and the Sadler-like effect vanishes under a constant rate of sequence evolution. The p-value associated with the hypothesis of zero slope is p = 0.04, up from p = 10^-15^ in the real dataset. The corresponding chronogram is shown in full on **Figure S16**.

## Discussion

### Tempo of molecular evolution and its alignment with morphological evolution

While the relationship between evolution at the molecular and morphological level is complex and non-linear, conceivably punctuations in molecular evolution could underlie punctuations in morphological change. The profound rate variability in molecular evolution across the evolutionary history of eukaryotes demonstrated here is consistent with this hypothesis. It is also noteworthy that we observe greatest departures from mean rates in parts of the eukaryotic tree best constrained by the fossil record – so it is likely that other, weakly constrained parts of the tree would also show profound substitution rate variability, if only fossil calibrations were available to better constrain divergence times there. To the extent that calibration densities admit such comparisons, periods of fast molecular evolution inferred here generally align with times of high evolutionary rates at the morphological level, such as close to the base of the animal tree in the Ediacaran and Cambrian (7), and with increases in morphological diversity, such as in early Mesozoic dinoflagellates (21, 22). While the limited resolution of fossil calibrations in our dataset only allows measurement of rate variability within molecular evolution over million-to billion-year timescales, it is likely that punctuations of smaller magnitude analogously occur on shorter timescales as well.

### Substitution rate variability and scale dependence in evolutionary interpretations

In addition to rate variability across the tree, we observe that inferred substitution rate tends to decrease with branch length across the whole dataset spanning million-to billion-year-long branches (**Figure 1** and **Figure 3**), This is beyond the threshold for the more commonly recognized reduction in rate between the appearance of mutations during short-term (generational) evolution on one hand and the fixation of substitutions at population/species scale on the other (e.g., (17)). Indeed, no stabilization of scale-dependent rates is evident even at a billion-year timescale, the longest available in the data. While prior literature has offered varying suggestions for causes of scale-dependence in different types of evolutionary rates (16, 23), we suggest – analogously to the widely accepted Sadler effect in sedimentary geology – that the rate variability present in this dataset of eukaryotic molecular substitution rates is responsible for scale-dependent behavior.

Accordingly, we demonstrate the absence of such scale-dependence when the same measurement technique is applied to a synthetic evolutionary history with sequences evolved at a constant rate (**Figure 4**). While mechanisms such as Brownian motion can generate scale-dependence even under constant rates (23), we thus do not identify this as a significant factor in long-term sequence evolution – perhaps because sequence space is very large in its number of dimensions and returning towards a prior state becomes vanishingly unlikely beyond a certain timescale.

While the exact cadence of rate variability is hard to pinpoint given the low resolution of available time calibrations, the variability should be such as to explain the Sadler-like effect: classically, longer periods of relative stasis should occur less frequently than shorter ones.

As for the selective mechanisms responsible, we have shown that they should apply genome-wide rather than driving the evolution of particular genes – at least in the case of Metazoa, where speed-ups and slowdowns in substitution rates from branch to branch are positively correlated between different genes and even the most clock-like genes considered show considerable rate variability (**Figure 2**). In future work, testing genes known to be under different selective pressures may illuminate the selective landscape responsible for the rate variability: while conserved proteins used for deep-time molecular clocks are often limited to slow time-averaged evolution by purifying selection, it is possible that it also creates punctuations. Proteins under positive selection – for example those involved in Metazoan immune responses (24, 25) – offer comparatively higher rates but potentially fewer punctuations. If they show reduced scale-dependence of substitution rate, that would further support the punctuation model of scale-dependence suggested here.

### Implications for molecular clock studies

Even if the sources of long-term substitution rate variability and scale-dependence are not entirely clear on the mechanistic level, these effects are profound enough to be considered in molecular clock models. Relaxed molecular clocks already allow some substitution rate variability, but the evidence considered here suggests that their bulk effect is to understate it in poorly calibrated clades. In explicitly quantifying extremes of rate variability in eukaryotic evolution, this study confirms the central role of calibrations for timing accuracy and precision in molecular clocks, even beyond the sequence data – in agreement with (26), and, in the context of early Metazoa specifically, with (27, 28). However, especially older-bound constraints are hard to come by even in clades with relatively better fossil records, and a literal interpretation of the absence of fossil evidence is challenging to justify (c.f., (29)).

A more serious omission in molecular clock studies is that substitution rates are not considered in the context of measurement timescale: in fact, we are not aware of any commonly used molecular clock approach that considers the scale-dependence of measurement in fitting substitution rates. In deep-time phylogenies such as considered here, branch lengths routinely cover multiple orders of magnitude, and we demonstrate that the consideration of the Sadler-like scale-dependence becomes essential. Furthermore, the magnitude of the Sadler-like effect shown in the posterior divergence time estimates is understated, as the clock model still pushes rates towards the same mean across all branch lengths. The clock runs performed only under the prior likely offer a better estimate of the magnitude of the Sadler effect. They show an approximately five-fold change in average substitution rates over two orders of magnitude in measurement timescale – a change so profound that it would clearly need to be considered by clock models, lest branch lengths be systematically misestimated by several times.

This effect is particularly concerning for short branches due to their typical importance in hypothesis testing. Wherever the interaction between evolutionary history and sampling choice breaks the tree up into short branches (such as close to the base of Metazoa), current molecular clocks overestimate branch lengths in time: not taking into account the Sadler-like effect, they underestimate substitution rates on those branches. Studies often naturally focus on resolving evolutionary histories and breaking up branches in parts of the tree most crucial to testing their hypotheses – and so the resulting time dilation is also most pronounced there, at least in the absence of tight fossil calibrations on these branches. As an example, molecular clock analyses have been applied to support an early Neoproterozoic origin of sponges (19, 27, 30, 31), 10s-100s of millions of years before the oldest uncontroversial sponge lipid biomarkers and body fossils appeared in the Ediacaran (32–36). The prevalence of the Sadler-like effect that we demonstrate here suggests such molecular clock-driven interpretations are prone to underestimate substitution rates on short branches around the base of Metazoa: the emergence of animals may not have occurred so early.

Even accepting that the crown nodes of Mollusca or Bilateria are well-constrained to the Ediaracan by the fossil record (calibrated with soft bounds spanning 17 Myr and 86 Myr, respectively, in (18)), working backwards using substitution rates to estimate the age of crown Metazoa (calibrated to a range covering ∼200 Myr from mid-Tonian to the beginning of the Ediacaran) or total group Metazoa (uncalibrated) is severely impacted by the Sadler-like effect. Calculated simply: starting from 550 Ma, by which time the fossil record contains uncontested examples of crown Bilateria (37), a realistic three-fold misestimation of sequence evolution rate due to omitting the Sadler-like effect is the difference between placing an ancestor node of interest (for example, the crown node of Metazoa) in Ediacaran at 600 Ma and placing it in early Cryogenian at 700 Ma. These results have profoundly different evolutionary implications, such as whether Metazoan clades survive multi-million-year Snowball Earth glaciations (38). A similar calculation could be repeated for whatever comparatively fixed point suggested by the fossil record. Together with the occurrence of actual rate variability (not simply scale-dependent changes in apparent substitution rate) in Metazoa, distinguishing between alternative evolutionary hypotheses in the case of animal origins becomes a daunting task for molecular clocks.

Beyond divergence times, estimates of past evolutionary rates – whether molecular or morphological – may not accurately reflect speed-ups or slow-downs if not considered in the context of measurement timescale. For example, Lee et al. (7) observe higher rates of morphological and molecular evolution in arthropods around the Cambrian Explosion than elsewhere on their tree, but these branches are also about an order of magnitude shorter than typical elsewhere on the tree. If the magnitude of the Sadler-like effect is comparable to our best estimate, this alone accounts for a two- to three-fold change in the inferred substitution rate, without any actual rate change. While this does not account for all of the observed speed-up and some rate change likely occurred, this explanation may also apply to a subsequent study on rates of morphological change in trilobites (8), which identified constant rates throughout the Cambrian – without their tree extending all the way to the present, branches across the tree are much more even in length and little Sadler-like effect is expected. Thus, for accurate estimation of both divergence times and evolutionary rates, models of evolution such as those used in molecular clocks should consider fitting branch rates in a scale-dependent manner. The strength of the scale-dependence could be a model input based on prior calibrations of the value, such as offered in this study, or it could be treated as a free parameter.

More broadly, we have argued that substitution rate variability and its scale dependence have been comparatively overlooked in molecular evolution and especially molecular clock studies, while being appreciated with more clarity in other fields such as sedimentary geology and suggested by punctuated dynamics long recognized in morphological evolution. Quantifying the profound degree of substitution rate variability and its scale dependence in a eukaryotic dataset, we outline its impact and suggest how it might be more explicitly recognized. This is crucial for the accurate interpretation of evolutionary history beyond what is apparent at face value in the fossil record. As with morphological evolution, an intriguing question is whether rate variability is predominantly internal to specific clades or branches, or the extent that it could be influenced by external, environmental factors – such as proposed changes in UV-B fluxes (39) associated with variation in Earth’s magnetic field intensity (40–42), extreme climate perturbations like Snowball-Hothouse intervals (38), or changes in surface oxygenation levels (43, 44). One path forward to consider this would be comparison of rates within disparately related parts of evolutionary trees to search for temporal correlation of intervals dominated by shorter branch lengths and faster substitution rates. For example, if the Ediacaran-Cambrian radiation of animals does indeed represent a time interval of enhanced rates of both morphological and molecular evolution (7), can we find evidence of correlative rate speed-ups in other distant eukaryotic or even bacterial clades? The challenge will continue to be the poor calibration of the non-metazoan record.

## Methods

We estimated substitution rate variability in datasets (sequence alignments and calibration schemes) used for two previously published molecular clock studies: (18) covering the evolutionary history of eukaryotes and (19) focusing on animals and close outgroups (including their supplemental data in (45)). We ran our main analyses on the dataset from (19) given its larger taxonomic breadth and higher number of time-constrained branches, allowing for more substitution rate measurements. We included (19) – providing a different taxonomic focus and including a different set of genes and fossil calibrations – to verify that our findings are not specific to a particular dataset.

Assembly of the datasets in (18, 19) has been described in the respective publications, with (19) further largely borrowing their dataset from (46). Each dataset was assembled to include a set of widely conserved single-copy genes showing congruent phylogenies and lack of evidence for horizontal gene transfer events.

### Substitution rate calculation

We measured the branch-specific substitution rates inferred on the autocorrelated relaxed clock (rooted on Amorphea) from (18) using their 136-taxon, 320-gene dataset (73,460 amino acid positions) and 33 fossil calibrations implemented as soft-bound uniform priors. We calculated substitution rates by dividing the length of each branch of the phylogenetic tree (reflecting the number of substitutions per site) with its length on the chronogram (reflecting the time between mean posterior ages of the nodes it connects) – see **Figure 1a**. We also calculated branch-specific substitution rates for the corresponding uncorrelated relaxed clock model to test the dependence of rates on clock model choice, and in an alternative dataset (19) to check that the observed patterns of rate variability are reproduced – see **Supporting Text**.

### Gene-by-gene rate variability

To understand the contribution of each gene to the aggregate rate variability signal, we examined the branch-dependent rates on a gene-by-gene basis within the subtree relating all Metazoa in the original dataset. This analysis was performed on 164 out of the original 320 genes, as each gene had to be present in every included Metazoan taxon for congruent branches to be available for comparison. We inferred maximum-likelihood gene trees in IQTree 3.0.1 (47) using a constraint tree that followed the topology of the chronogram described above. We used the LG substitution matrix (48) with equilibrium frequencies estimated from the data and four gamma-distributed site rate categories.

We then calculated branch-specific substitution rates for each gene by dividing the length of each branch of the gene tree with its length on the chronogram (between mean posterior node ages). We then examined rate variability from branch to branch by creating a histogram of branch rates normalized to mean rate for the gene and calculating the standard deviation of that distribution for each gene (**Figure 2a**). We also performed a pairwise regression between rates on corresponding branches for each pair of genes using the linregress function in the SciPy Statistics library and recorded the R-value for each pair (**Figure 2b)**.

### Testing the scale-dependence in substitution rate estimates

To quantify any scale-dependent observation bias in substitution rates, we plotted on a log-log scale the substitution rate inferred on each branch against the branch length between mean posterior node ages. We then performed a linear regression between these variables using the polyfit function in Python’s NumPy library and recorded the slope of the regression line as a proxy for the timescale dependence. Given the log-log scale, the units of slope value show orders of magnitude of change in inferred substitution rates per order of magnitude of change in branch length. We tested the statistical significance of the non-zero slope with the linregress function in the SciPy Statistics library.

### Testing the impacts of sequence data, internal node calibrations, and tree prior

To understand the impact of different components of the clock on the inferred scale-dependent observation bias, we re-analyzed the dataset from (18) described above, making relaxed molecular clocks under different combinations of 1) including or omitting sequence data (running the clock under the prior only), 2) including or omitting fossil calibrations on internal nodes, and 3) tree prior choice (birth-death or uniform) – see **Table 1**. For computational efficiency and to ensure good mixing of the chains, we performed these analyses on a subsample (without replacement) of 1000 sites from the original alignment, obtained using a custom Python script implementing BioPython tools. Where internal calibrations were included, we used the same set of 33 calibrations as (18), applied as uniform priors with hard bounds.

We performed the relaxed molecular clock runs with PhyloBayes 4.1 (49–51), using the autocorrelated rate model with rates drawn from a log-normal distribution (“-ln”). We used the LG substitution matrix (48) with equilibrium frequencies estimated from the data and four gamma-distributed site rate categories. We required two chains to converge before sampling the posterior distributions. In accordance with recommendations in the PhyloBayes manual, we assessed convergence by requiring TRACECOMP values for all estimated variables to be <0.3 and the minimum effective size of the sample to be >50. A 20% burn-in was used for all posterior samples and when checking for convergence. For runs with no sequence data included, the clock was run under the prior only (“-prior”), and for runs with no internal calibrations, the calibration file was modified to include only the root prior (uniform between 1.6 and 3.2 Ga, as in (18)).

All custom Python scripts used for data processing and visualization are included in the supplemental data files. The scripts were written with the aid of the GPT-4 large language model, with the initial generative output manually tested and modified.

### Simulated sequence alignment

To test our inference of rate variability on a negative control, we evolved a synthetic multiple sequence alignment 1000 sites in length along the branches of the autocorrelated molecular clock inferred above (the version including both internal calibrations and sequence data, and a uniform tree prior), scaled by the average substitution rate inferred for that clock (0.242 substitutions per site per Ga). This simulates an alignment evolved at a constant substitution rate across all the branches of the whole tree. We used the “--alisim” option in IQTree 3.0.1 (47, 52) with the LG substitution model. We then ran a molecular clock analysis on this synthetic alignment, using PhyloBayes 4.1 as above with the LG substitution model, equilibrium frequencies estimated from the data, four gamma-distributed site rate categories, the tree topology used to create the synthetic alignment, and no internal calibrations.

## Data availability

The supplemental data files for this manuscript, including Python scripts written for data analysis and visualization, as well as all alignment, tree, and chronogram files, can be accessed on Figshare at 10.6084/m9.figshare.32330592.

## Acknowledgments

We would like to thank authors whose data was used in this study for making their alignments, trees, and metadata publicly available, and we would additionally like to thank Fabien Burki for sharing the phylogenetic tree used for molecular clocks in (18). We acknowledge funding from the Department of Earth, Atmospheric & Planetary Sciences at the Massachusetts Institute of Technology and from the William H. Neukom 1964 Institute for Computational Science at Dartmouth College. E.T. is thankful to Greg Fournier for advice and access to computational resources.

## Notes

https://doi.org/10.6084/m9.figshare.32330592

### Competing Interest Statement

The authors have declared no competing interest.

