## Supplemental Information for "Variability and scale-dependence in molecular evolution rate impacts interpretation of eukaryotic evolutionary histories"

Erik Tamre

**This PDF file includes:**

Supporting text

Figures S1 to S24

SI References

Supporting Information Text

**Substitution rate variability and scale-dependence inferred with an uncorrelated molecular clock model**

To check if inferred substitution rate variability or scale-dependence significantly changes as a function of clock model used, we analyzed substitution rates inferred in (1) using an uncorrelated relaxed clock model – in addition to the autocorrelated (log-normal) clock model considered in the main text. In an uncorrelated clock model, there is no expectation of similar substitution rates on adjacent branches, and the inference of significantly different rates on adjacent branches is not penalized. As a result, the chronogram using the uncorrelated clock model (**Figure S17**) shows more rate changes from branch to branch, but general patterns discussed in the main text are still present: for example, significantly higher rates close to the base of Metazoa than in crown vertebrates, and elevated rates within dinoflagellates after the divergence of Noctilucales. The strength of recovered scale-dependence is similar to the autocorrelated clock model, with the slope of the log-log plot of inferred rate vs. branch length (**Figure S18**) slightly decreased to -0.19 (-0.21 for the autocorrelated clock).

**Substitution rate variability and scale-dependence in an additional dataset focused on Metazoa**

To check that the observed patterns of substitution rate variability and scale-dependence are not specific to the dataset in (1), we also studied substitution rates in another dataset (2) focusing specifically on animals and close outgroups. We calculated inferred substitution rates for the authors’ preferred fossil calibration set A and autocorrelated (log-normal) relaxed clock model, using their 1000 Ma root age analysis and testing both uniform and birth-death tree priors. To check for dependence of substitution rate patterns on root age, we also calculated substitution rates for the clock using an 800 Ma root age with the uniform tree prior.

Full chronograms showing rates from these analyses are displayed on **Figure S19-S21**, and the corresponding log-log plots of inferred rate vs. branch length on **Figure S22-S24**. Even though this dataset has a different taxonomic focus (with much finer sampling among Metazoa) and includes a different set of genes (making comparisons of absolute substitution rates in the two datasets not meaningful), the general patterns of substitution rate variability are similar within clades included in both datasets. Particularly high rates are recovered close to the base of Metazoa, and a slightly stronger Sadler-like scale-dependence effect is present than in analyses based on (1), with the slope of the log-log plot of inferred rate vs. branch length falling between -0.29 and -0.31 in all tested clocks from (2). It is likely that a more significant scale-dependence is recovered in this dataset because its taxonomic focus on Metazoa and close relatives has allowed more branches to be well-constrained by the fossil record, leaving less room for the clock model to artificially draw rates towards the mean. Tighter constraints on internal node ages may also explain why root age (mainly impacting comparatively poorly calibrated early nodes) and tree process prior choice have limited impact on the outcomes.

**Strength of scale-dependence in the original clocks from (1) and re-runs in this study**

The degree of scale-dependence recovered in our analysis of the original auto-correlated relaxed molecular clock from (1), described on **Figure 1**, is slightly lower than in the corresponding re-run (**Figure 3a**) performed as part of analyzing impacts different components of the molecular clock have on rate variability: the slope of the log-log plot of inferred rate vs. branch length is -0.214 and -0.26, respectively. While we used different phylogenetic software and performed the analysis on a subsampled sequence dataset (increasing the sample size from 1000 to 5000 sites was tested, but did not yield significant differences in rate estimates), the most important difference is probably that we implemented the fossil-calibrated node age constraints with hard bounds – rather than soft bounds, as in (1), with 2.5% of the probability mass falling outside the calibration range on each end. This choice reflects the suggestion from the original molecular clock that fossil calibrations are the most important source of timing information, and they should be leveraged without allowing the clock model to drive rates towards the mean excessively. That said, the choice of soft vs. hard bounds has the most impact where sequence data and fossil constraints are in conflict, so this choice does not significantly impact prior-only runs which we interpret as offering the best-guess estimate of the degree of scale-dependence in this dataset.

Figures

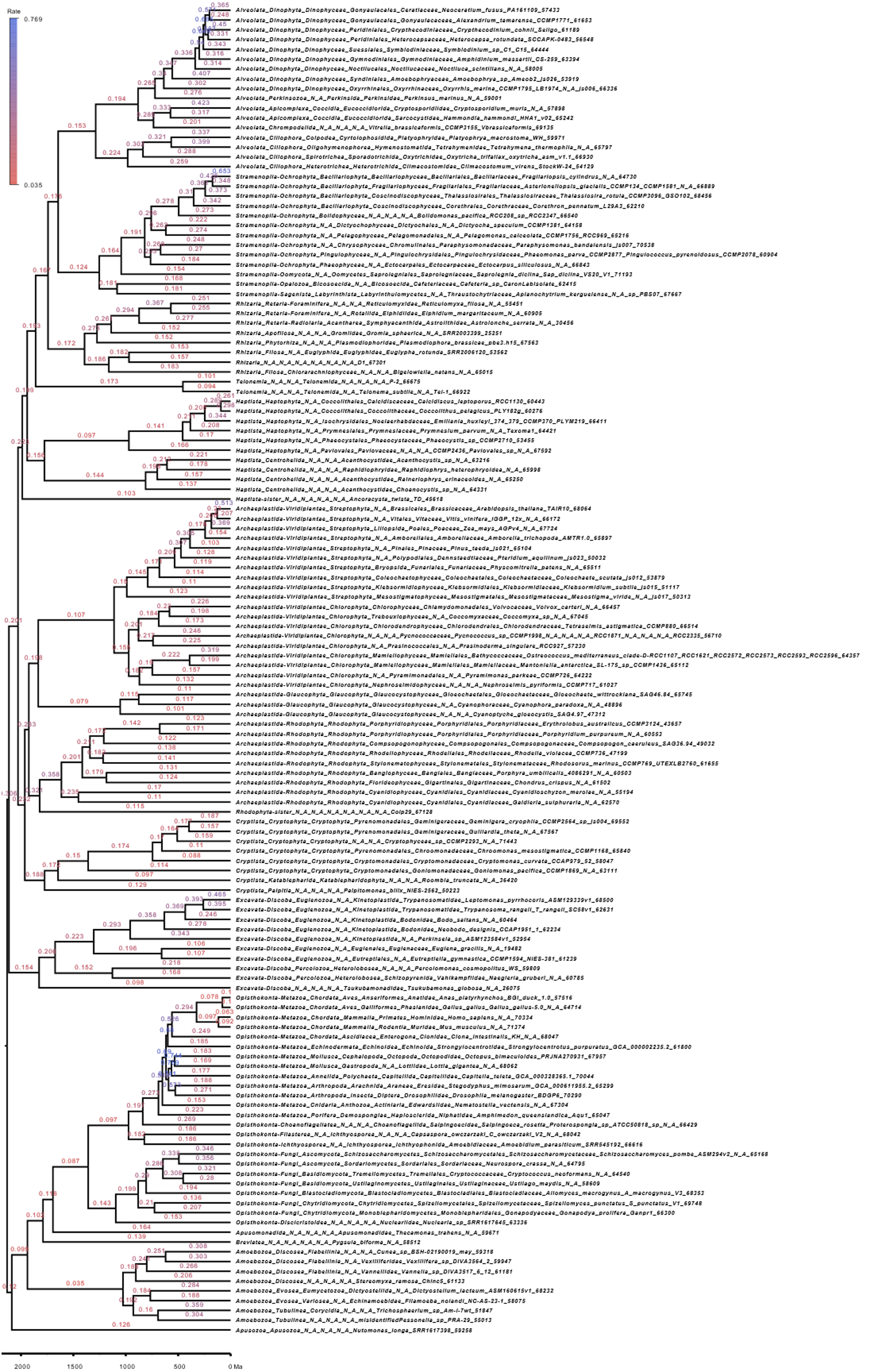

Fig. S1. Full chronogram showing inferred substitution rates in the original analysis in (1). Branch labels show the average substitution rate (in substitutions per site per Gyr) inferred along each branch using an autocorrelated relaxed clock model, with the tree rooted on Amorphea.

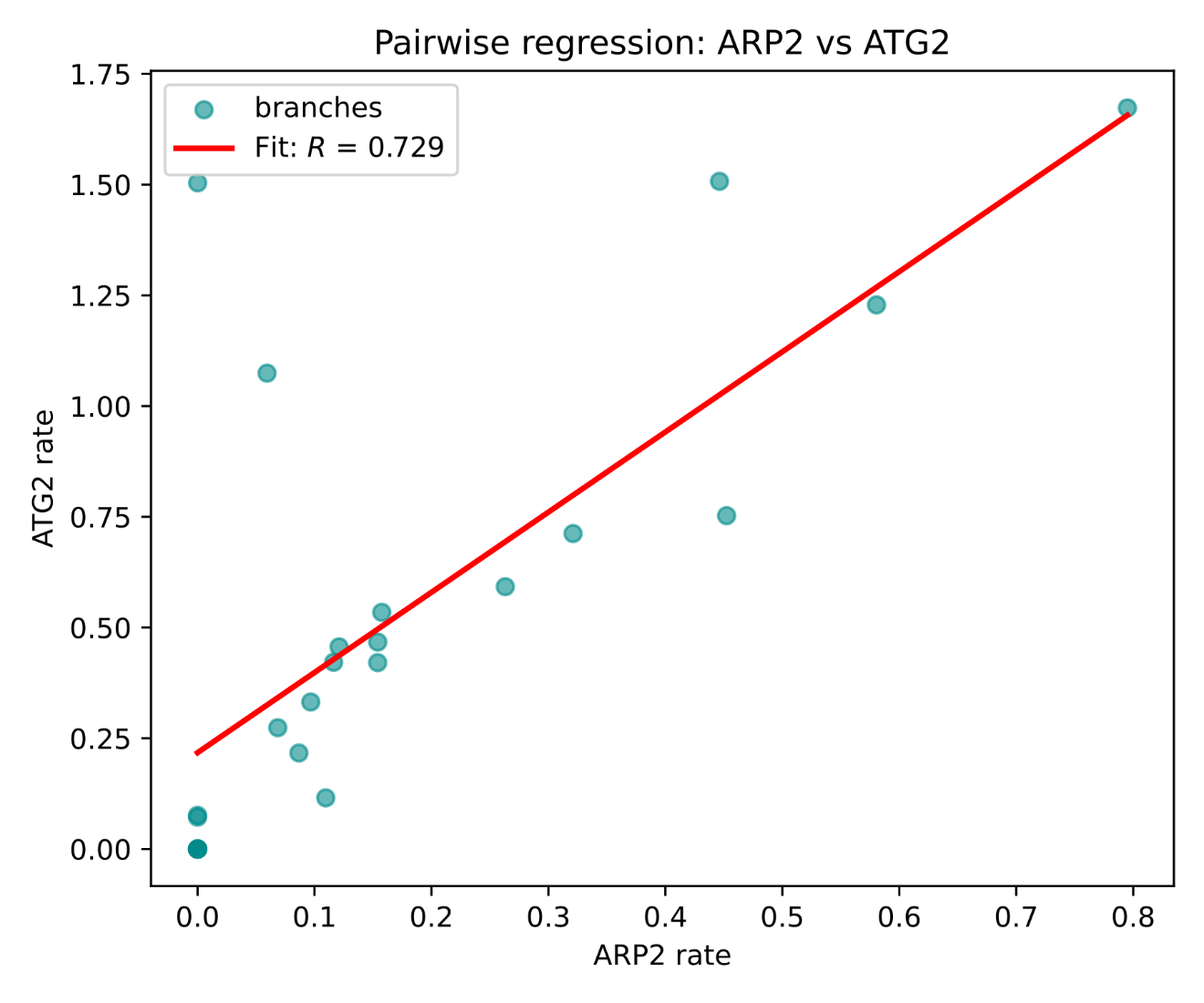

Fig. S2. Example of a histogram showing R-values for a regression between rates on corresponding branches in the metazoan part of the tree for one pair of genes: ARP2 and ATG2. Note that multiple branches plot to the origin, corresponding to no changes in either of the genes along a given branch.

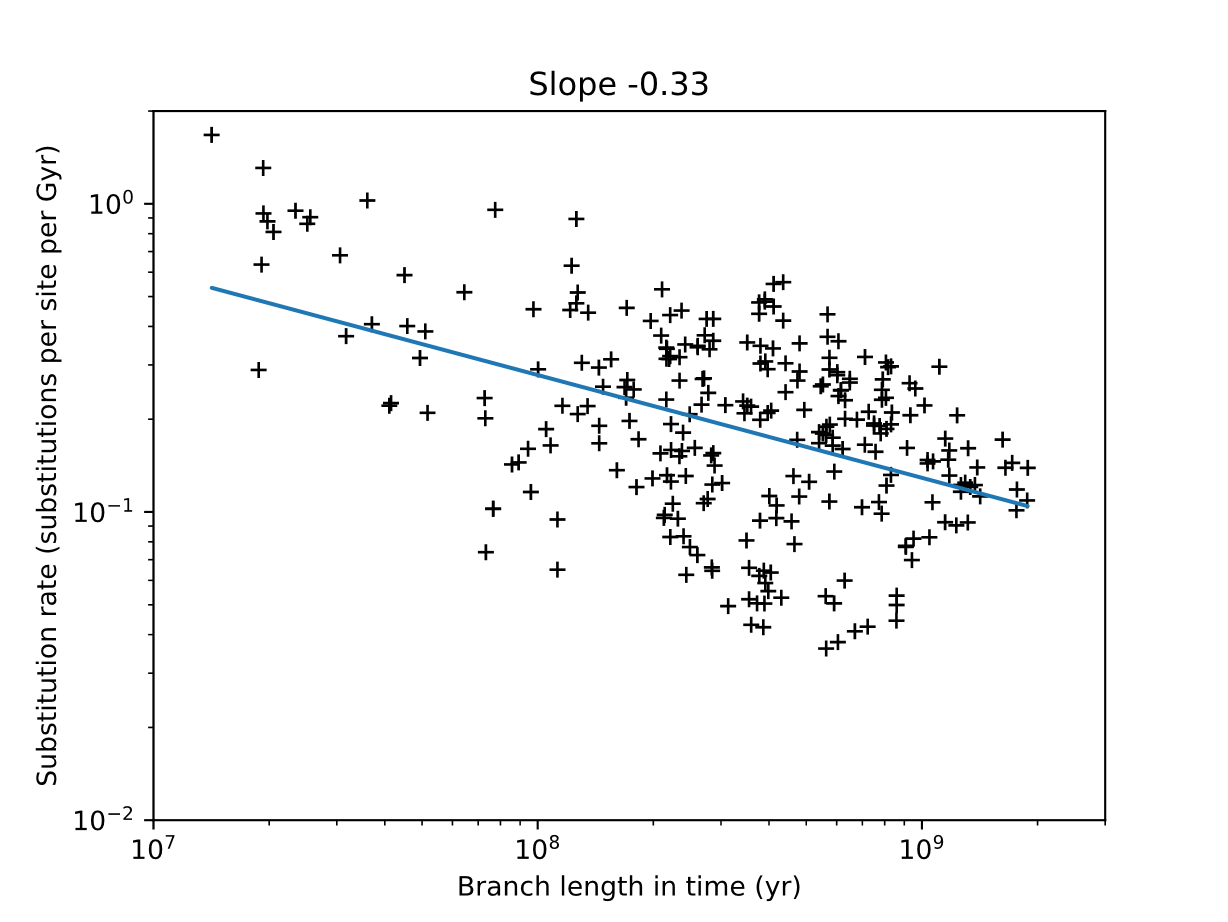

Fig. S3. The Sadler-like scale dependence of substitution rates in the dataset from (1) in a prior-only molecular clock run (no sequence data included) under a uniform tree prior, with internal calibrations included. The corresponding chronogram is shown in full on Figure S12.

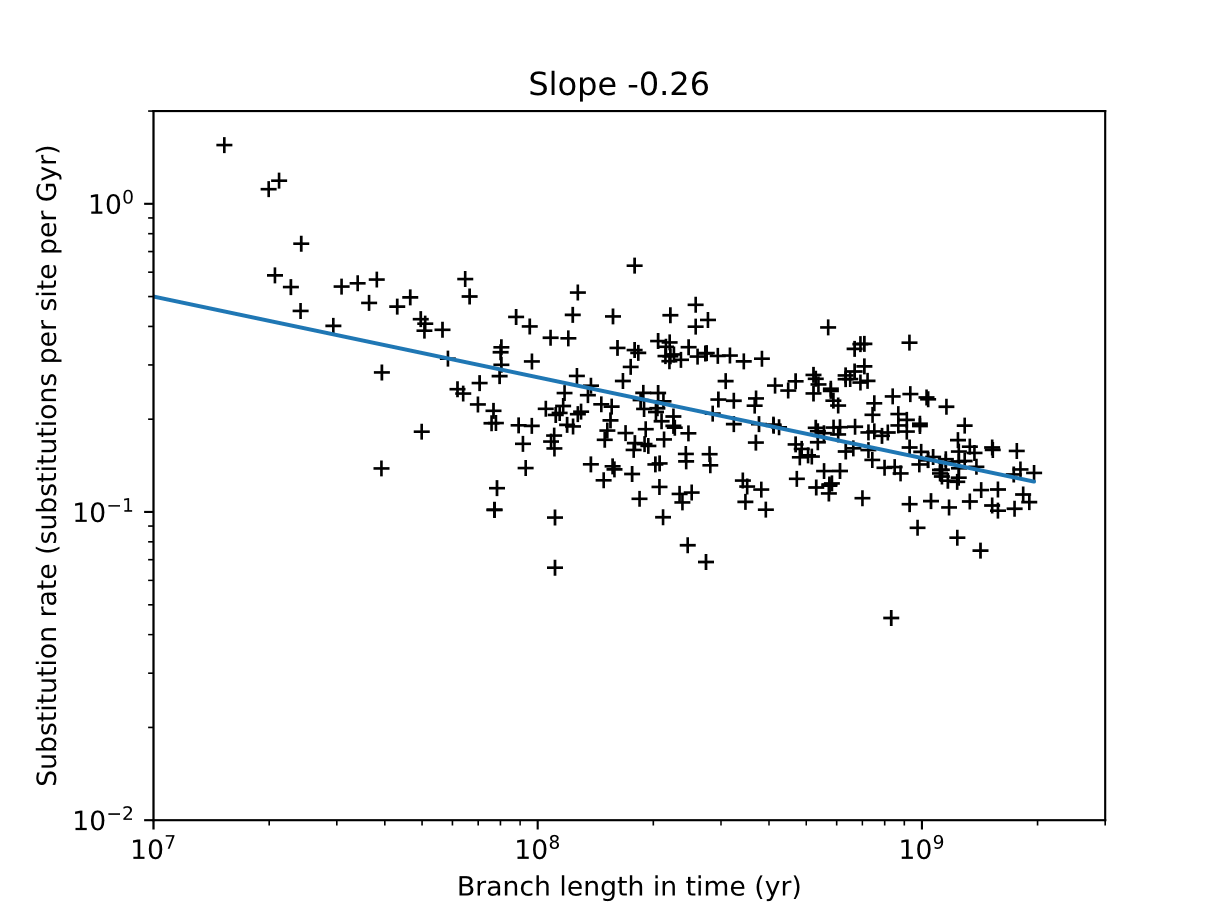

Fig. S4. The Sadler-like scale dependence of substitution rates in the dataset from (1) in a molecular clock run including the sequence data with an autocorrelated (log-normal) clock model and a uniform tree prior, with internal calibrations included. The corresponding chronogram is shown in full on Figure S13.

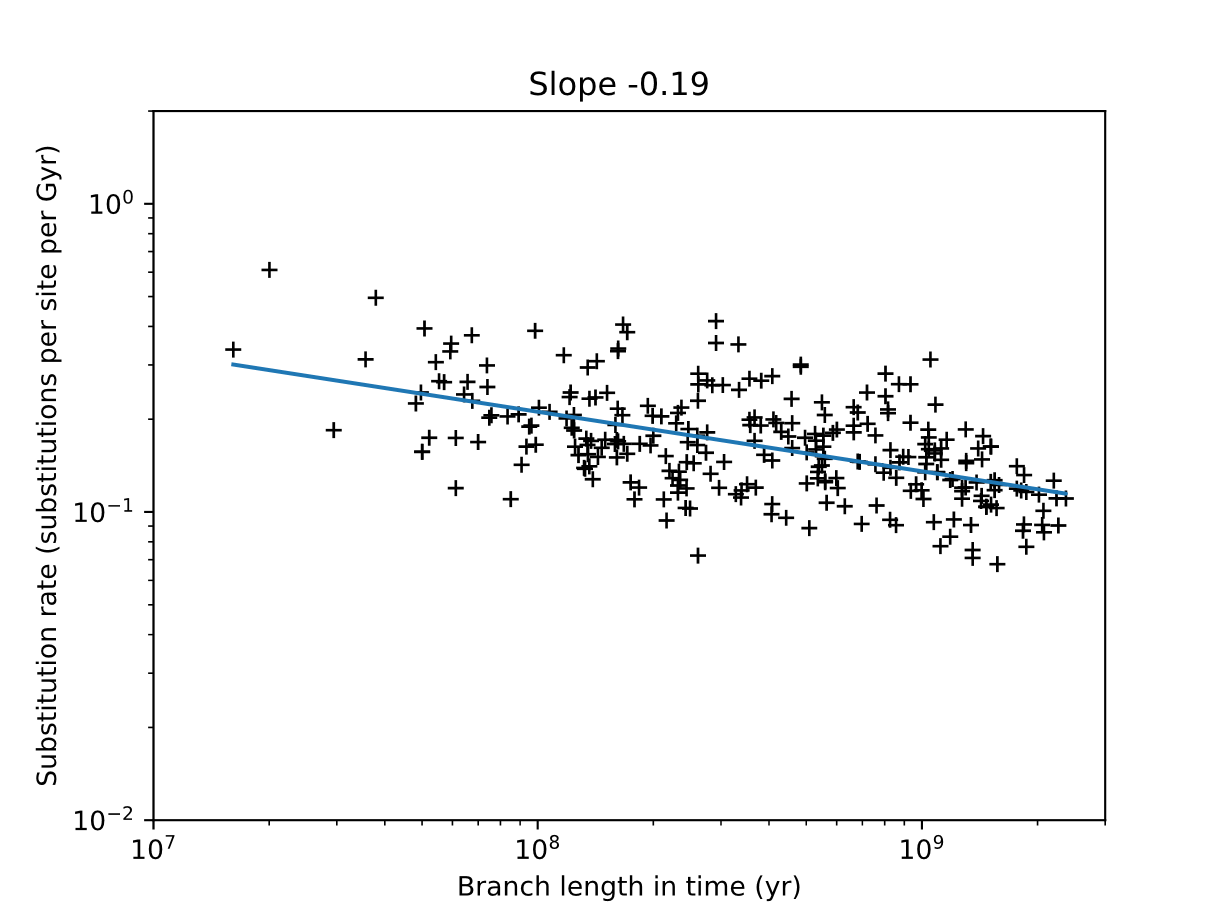

Fig. S5. The Sadler-like scale dependence of substitution rates in the dataset from (1) in a molecular clock run under a uniform tree prior, including sequence data and using an autocorrelated (log-normal) clock model, but omitting internal calibrations. The corresponding chronogram is shown in full on Figure S14.

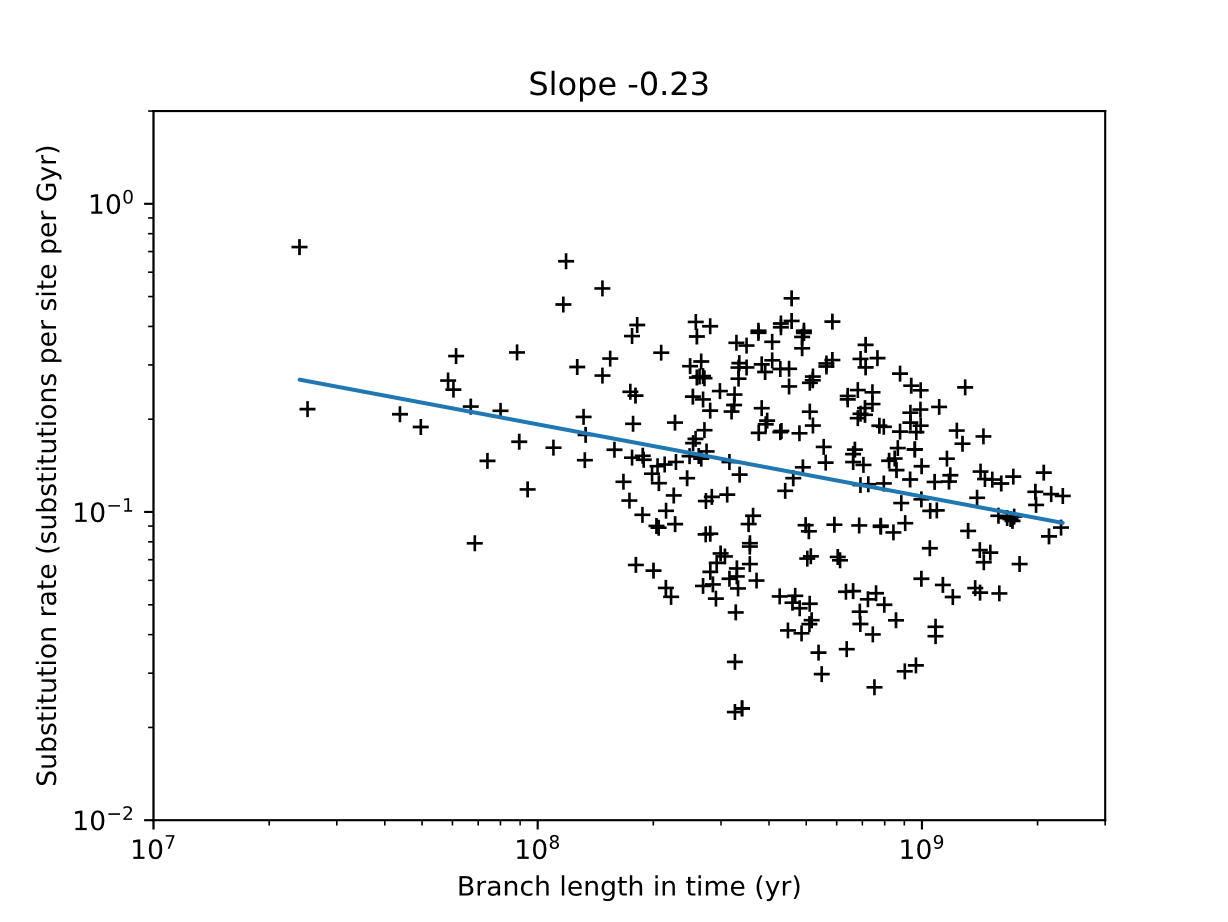

Fig. S6. The Sadler-like scale dependence of substitution rates in the dataset from (1) in a molecular clock run under a uniform tree prior, omitting both sequence data and internal calibrations. The corresponding chronogram is shown in full on Figure S15.

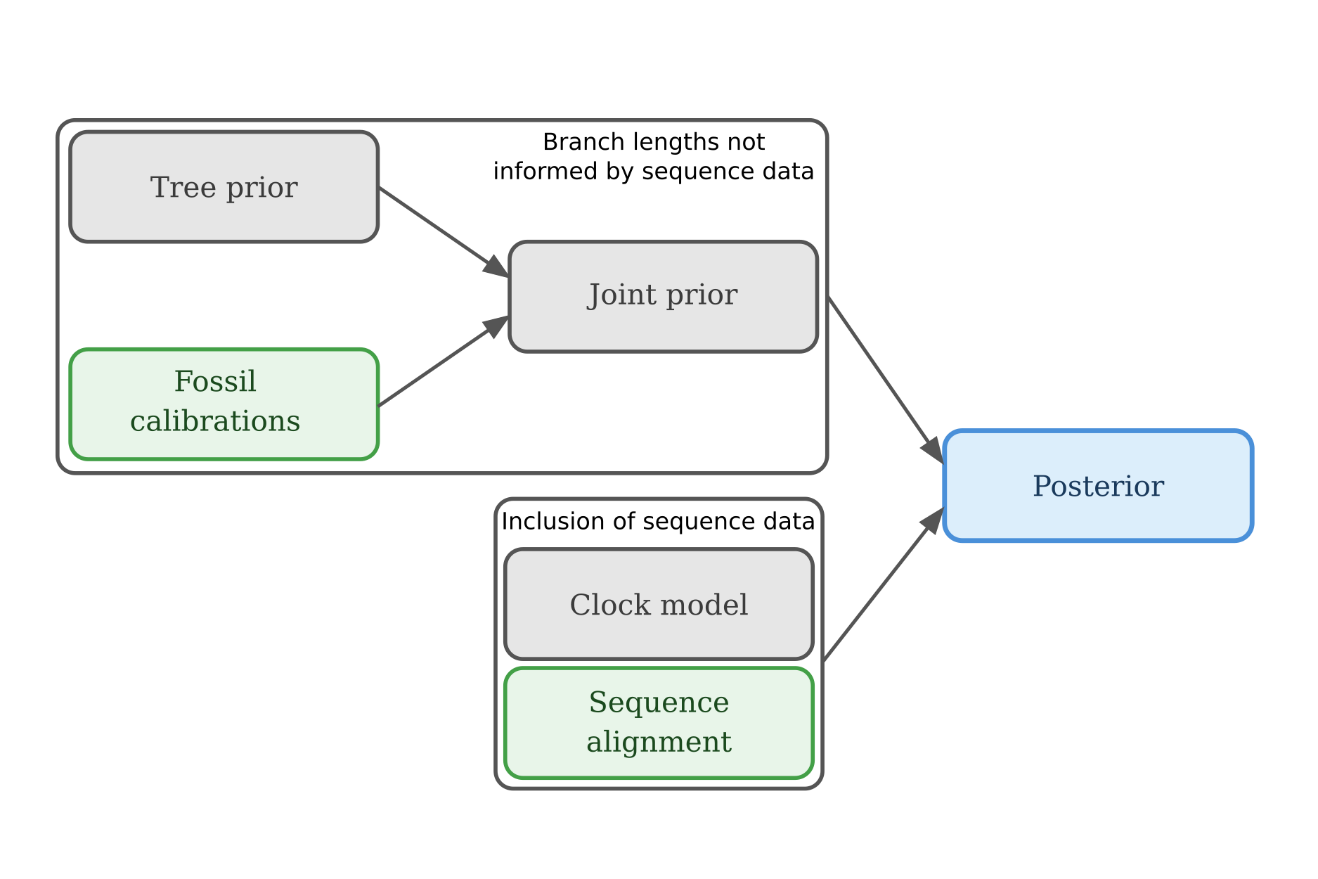

**Fig. S7.** A simplified schematic of the Bayesian molecular clock workflow. Note that while the fossil calibration data is included at the prior stage, sequence data is only incorporated upon the transition from prior to posterior.

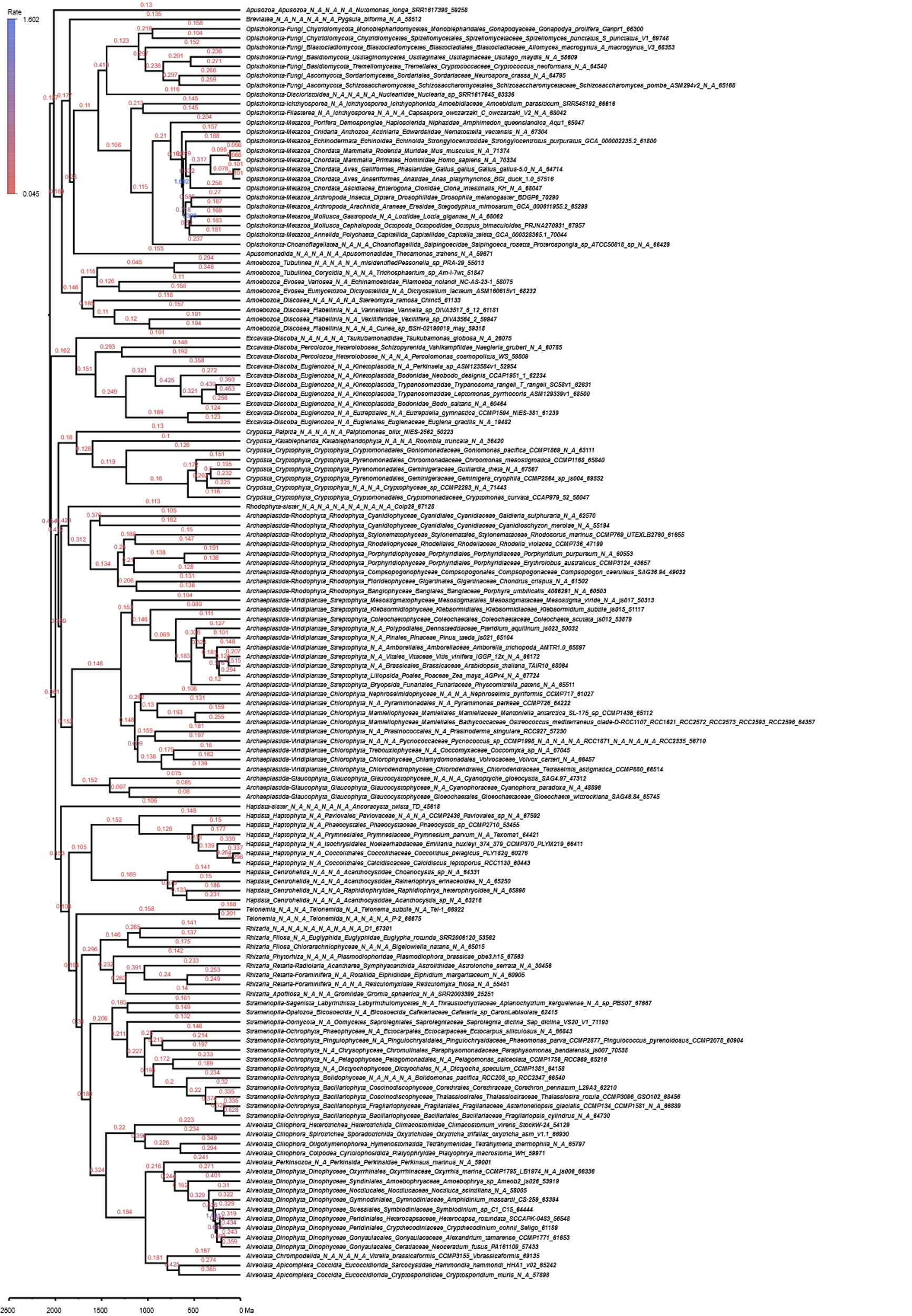

**Fig. S8.** Full chronogram showing inferred substitution rates in the analysis depicted on **Figure 3a**, using an autocorrelated (log-normal) relaxed clock model with a birth-death tree prior and including both internal calibrations and sequence data. Branch labels show the average substitution rate (in substitutions per site per Gyr) inferred along each branch.

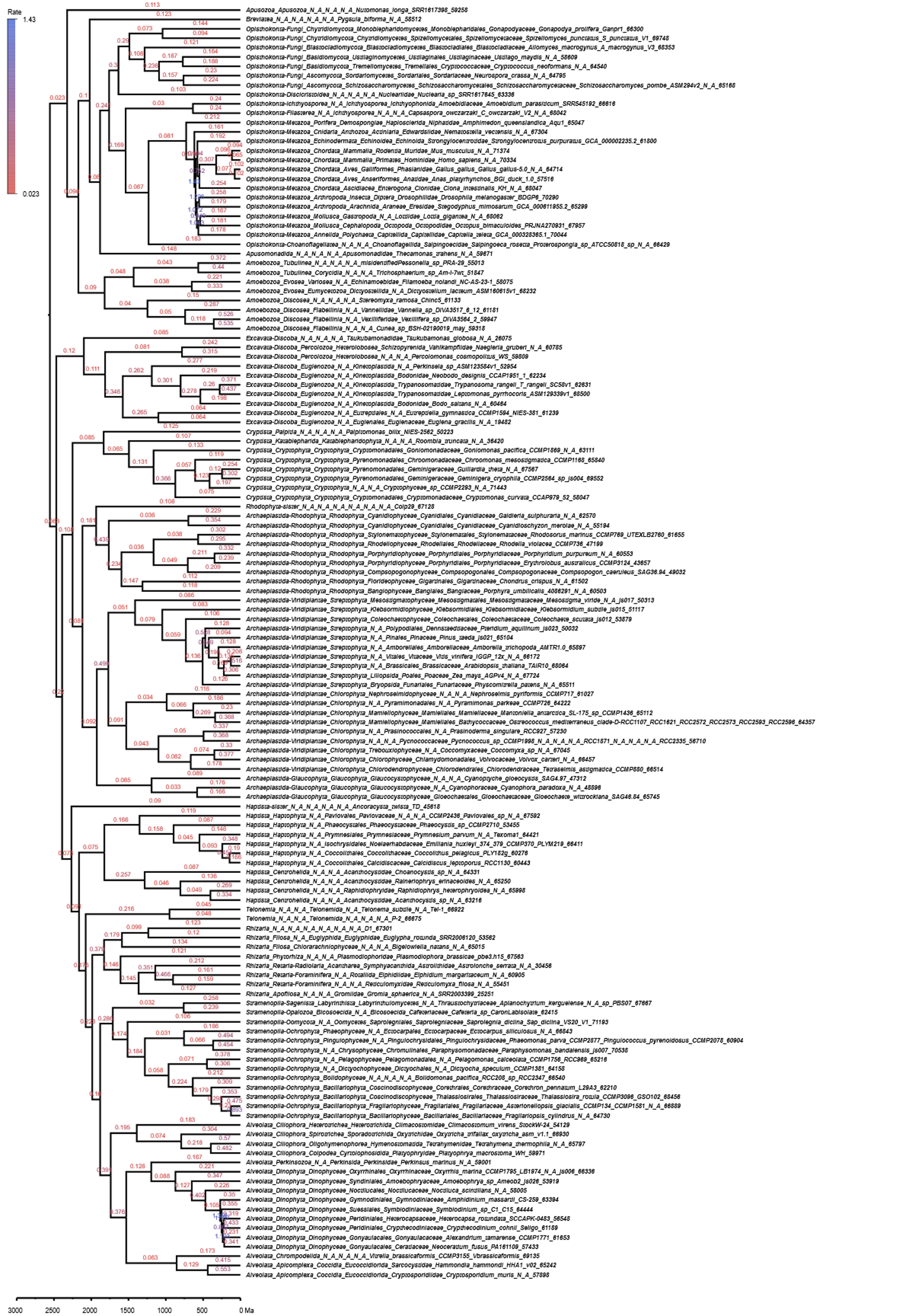

**Fig. S9.** Full chronogram showing inferred substitution rates in the analysis depicted on **Figure 3b**, using a birth-death tree prior and including internal calibrations, but no sequence data. Branch labels show the average substitution rate (in substitutions per site per Gyr) inferred along each branch.

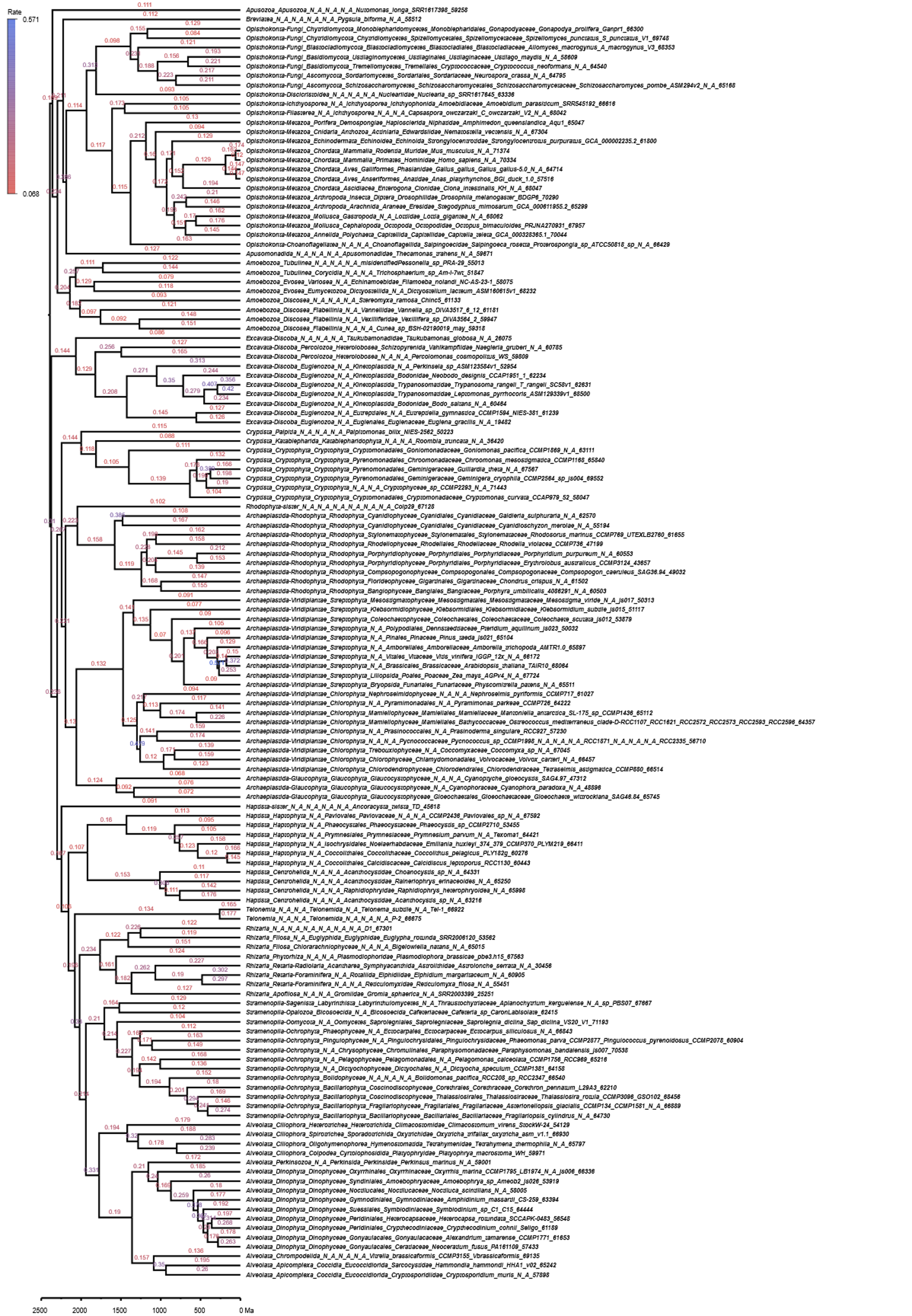

**Fig. S10.** Full chronogram showing inferred substitution rates in the analysis depicted on **Figure 3c**, using an autocorrelated (log-normal) relaxed clock model with a birth-death tree prior, but not including internal calibrations. Branch labels show the average substitution rate (in substitutions per site per Gyr) inferred along each branch.

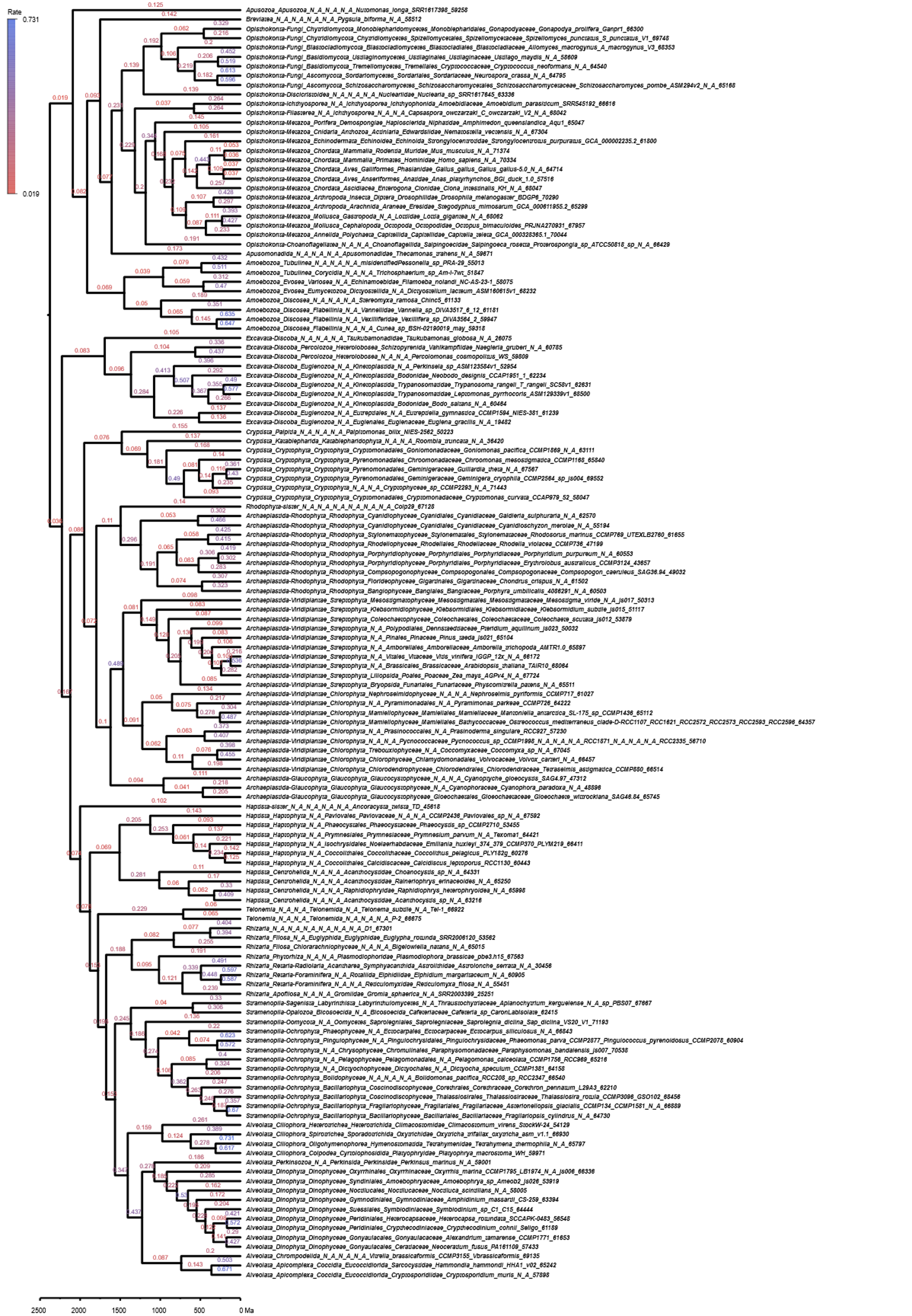

**Fig. S11.** Full chronogram showing inferred substitution rates in the analysis depicted on **Figure 3d**, using a birth-death tree prior, but not including sequence data or internal calibrations. Branch labels show the average substitution rate (in substitutions per site per Gyr) inferred along each branch.
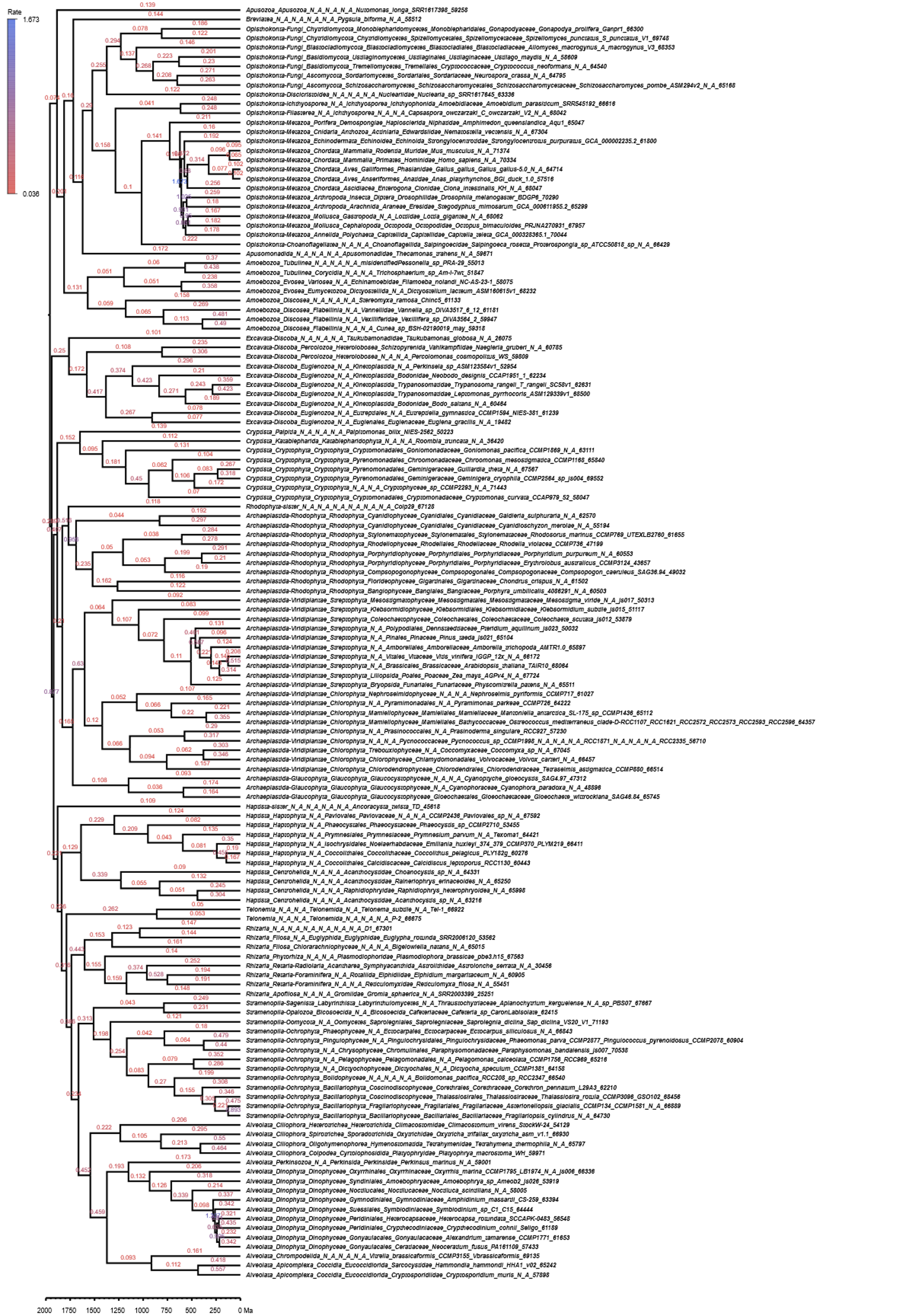

**Fig. S12.** Full chronogram showing inferred substitution rates in the analysis depicted on **Figure S3**, using a uniform tree prior and including internal calibrations, but no sequence data. Branch labels show the average substitution rate (in substitutions per site per Gyr) inferred along each branch.

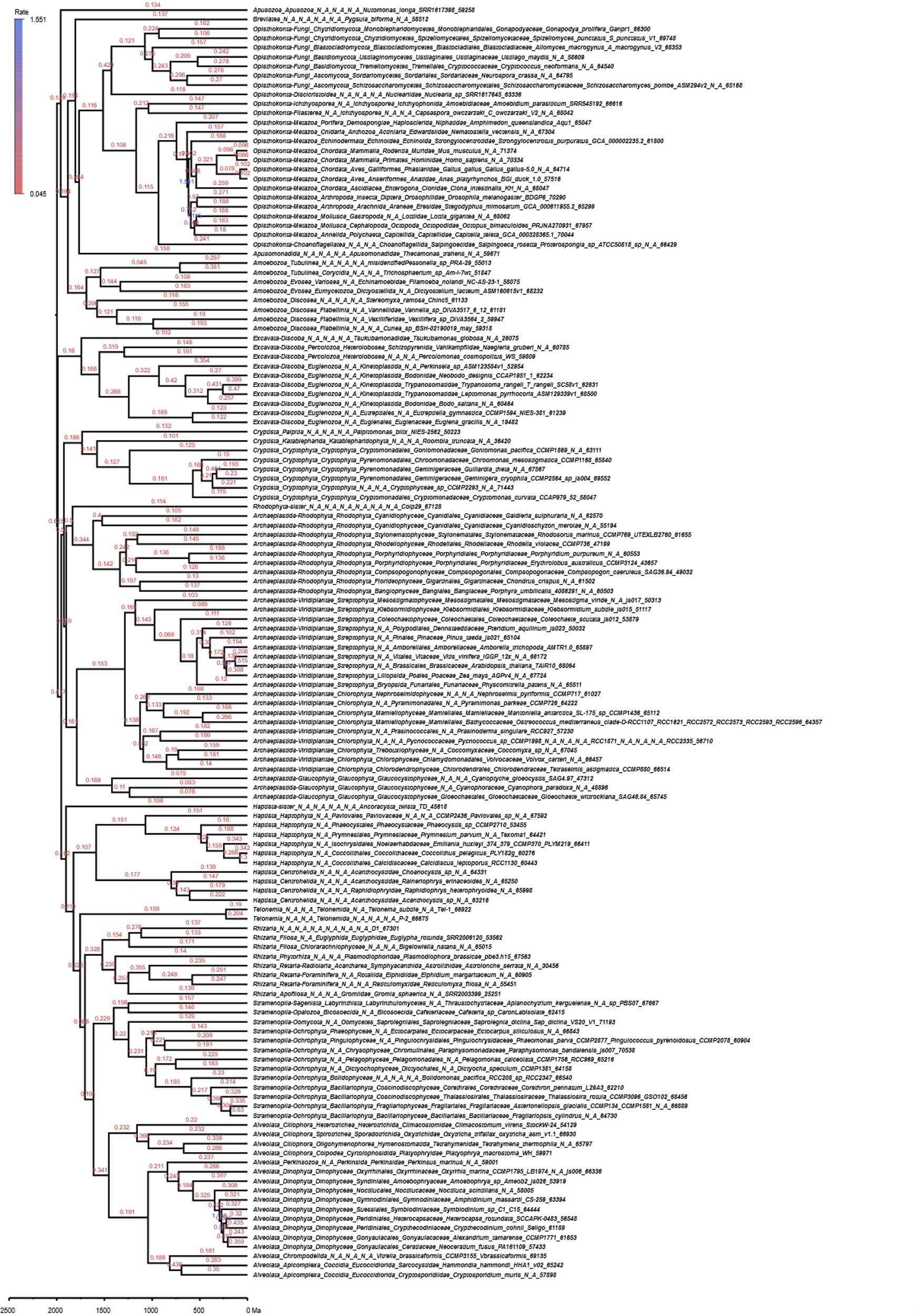

**Fig. S13.** Full chronogram showing inferred substitution rates in the analysis depicted on **Figure S4**, using an autocorrelated (log-normal) relaxed clock model with a uniform tree prior and including both internal calibrations and sequence data. Branch labels show the average substitution rate (in substitutions per site per Gyr) inferred along each branch.

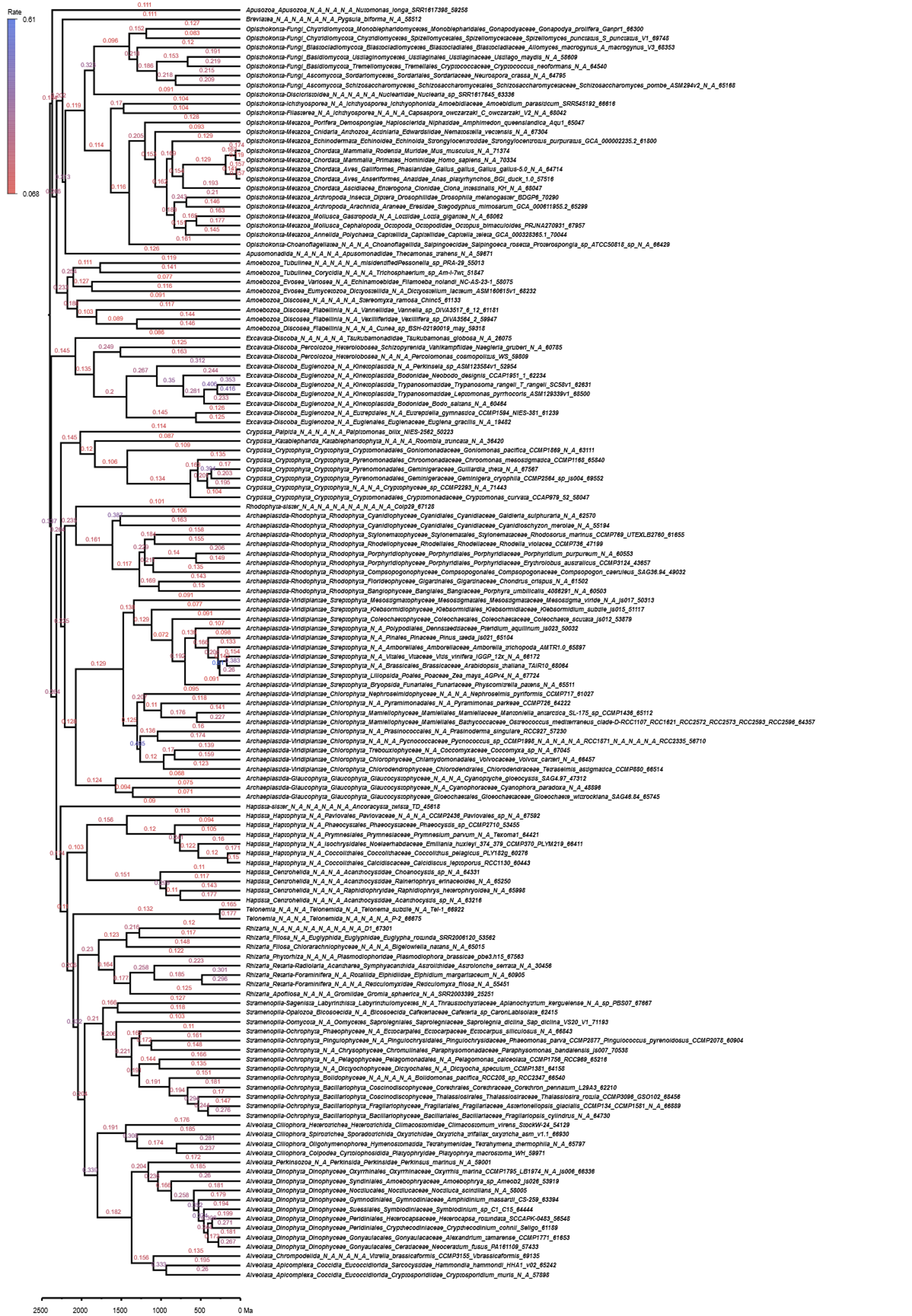

**Fig. S14.** Full chronogram showing inferred substitution rates in the analysis depicted on **Figure S5**, using an autocorrelated (log-normal) relaxed clock model with a uniform tree prior, but not including internal calibrations. Branch labels show the average substitution rate (in substitutions per site per Gyr) inferred along each branch.

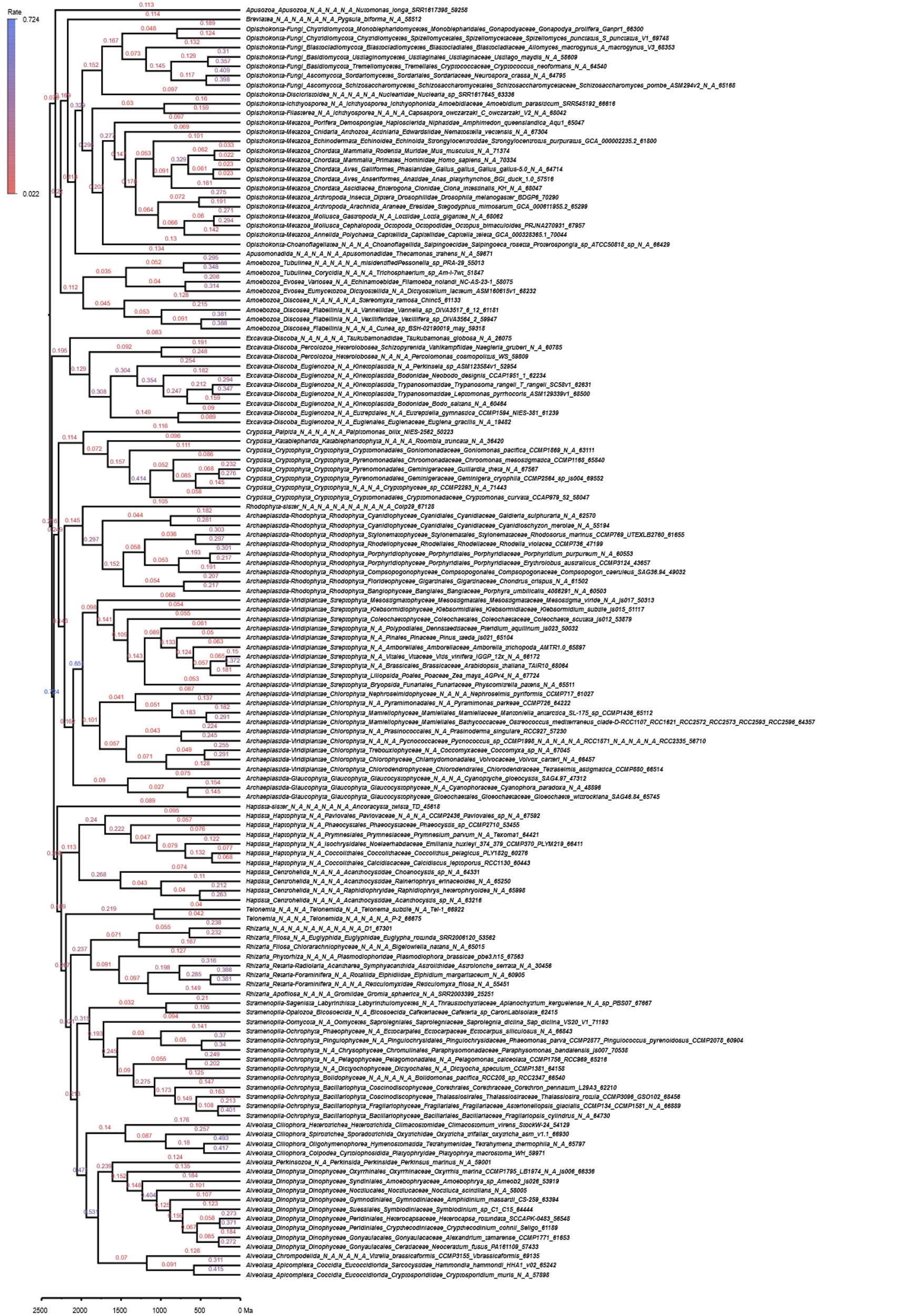

**Fig. S15.** Full chronogram showing inferred substitution rates in the analysis depicted on **Figure S6**, using a uniform tree prior, but not including sequence data or internal calibrations. Branch labels show the average substitution rate (in substitutions per site per Gyr) inferred along each branch.
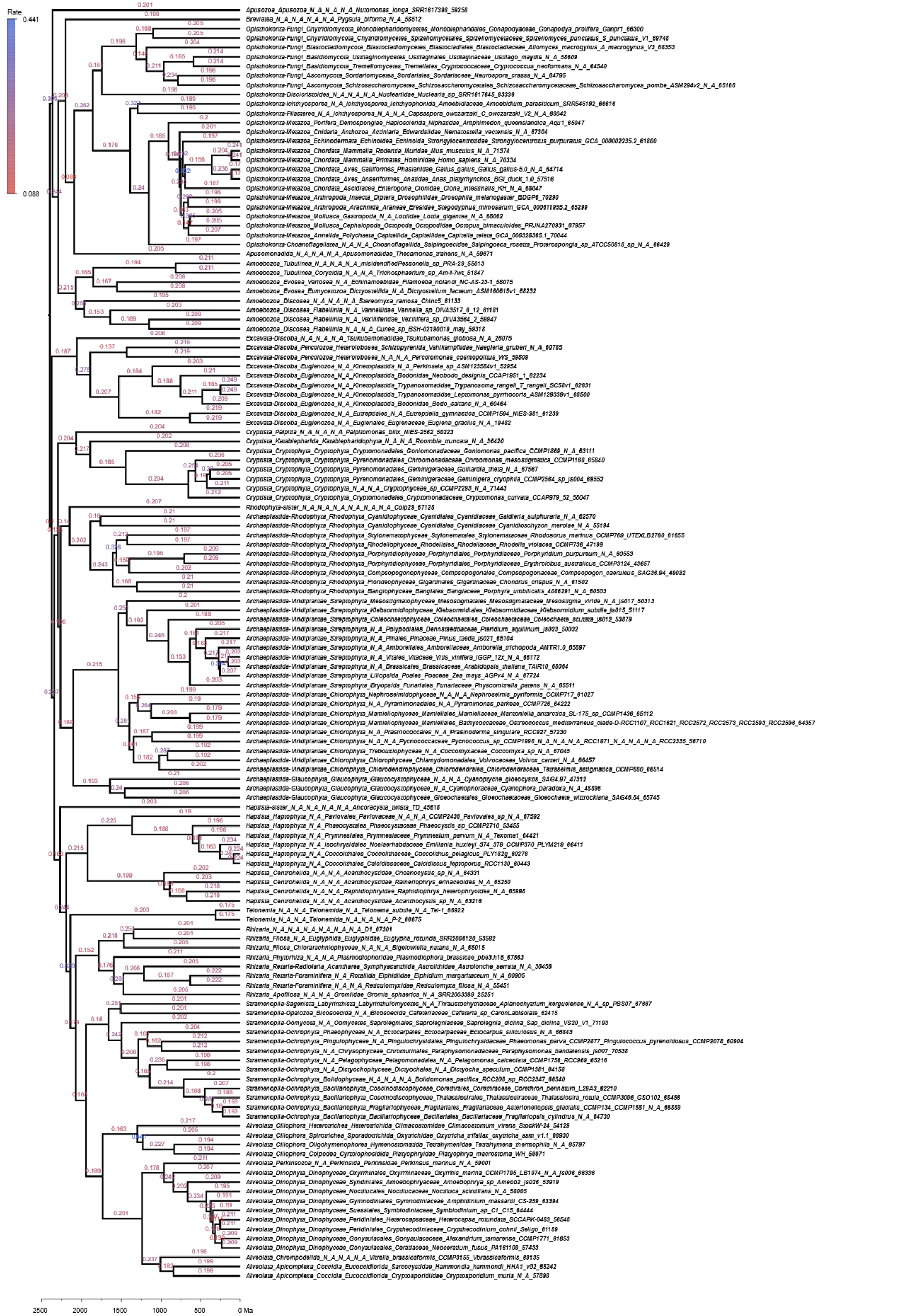

**Fig. S16.** Full chronogram showing inferred substitution rates in the analysis depicted on **Figure 4**, using a synthetic sequence dataset evolved at a constant substitution rate along branches of the molecular clock on **Figure 1**. Branch labels show the average substitution rate (in substitutions per site per Gyr) inferred along each branch.

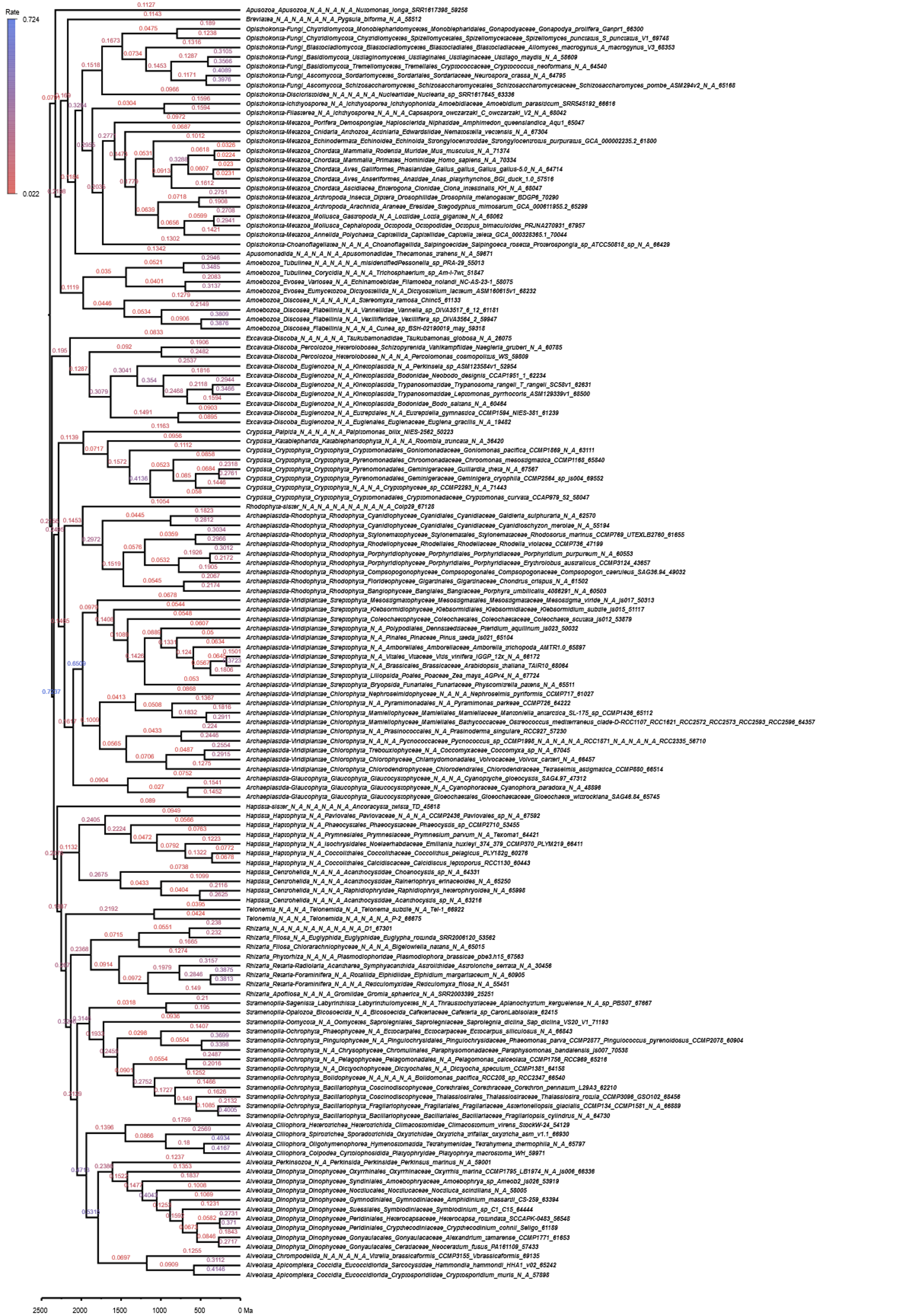

**Fig. S17.** Full chronogram showing inferred substitution rates in the original analysis in (1), using an uncorrelated relaxed clock model, with the tree rooted on Amorphea. Branch labels show the average substitution rate (in substitutions per site per Gyr) inferred along each branch.

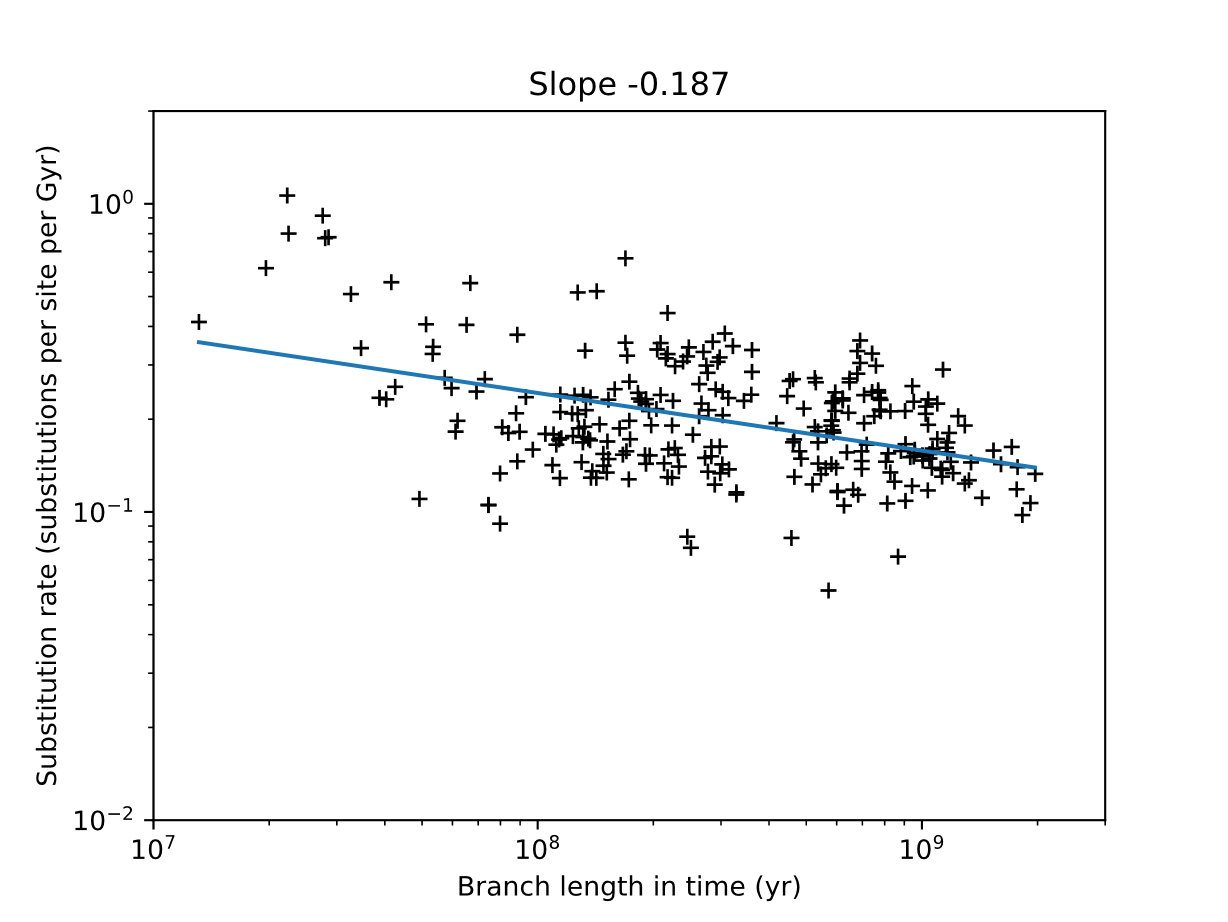

Fig. S18. The Sadler-like scale dependence of substitution rates in the dataset from (1) in a molecular clock using an uncorrelated clock model with branch rates drawn from a gamma distribution (as opposed to Figure 1, with an autocorrelated clock model). The corresponding chronogram is shown in full on Figure S17.

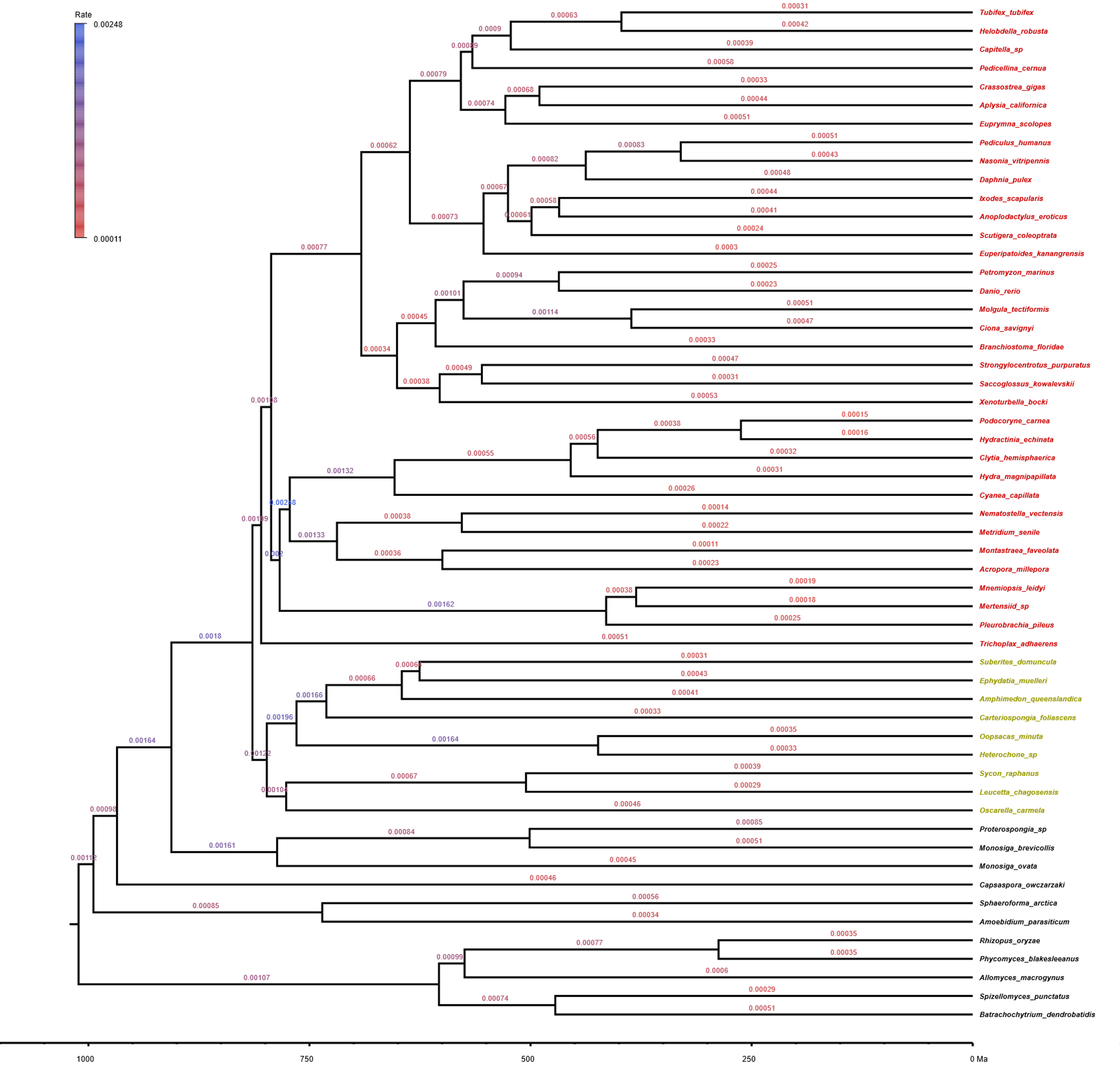

**Fig. S19.** Full chronogram showing inferred substitution rates in an autocorrelated (log-normal) relaxed molecular clock from (2) with a uniform tree prior and root age of 1000 Ma. Branch labels show the average substitution rate (in substitutions per site per Myr) inferred along each branch. Sponges are shown in yellow, other Metazoa in red, and non-Metazoa in black.

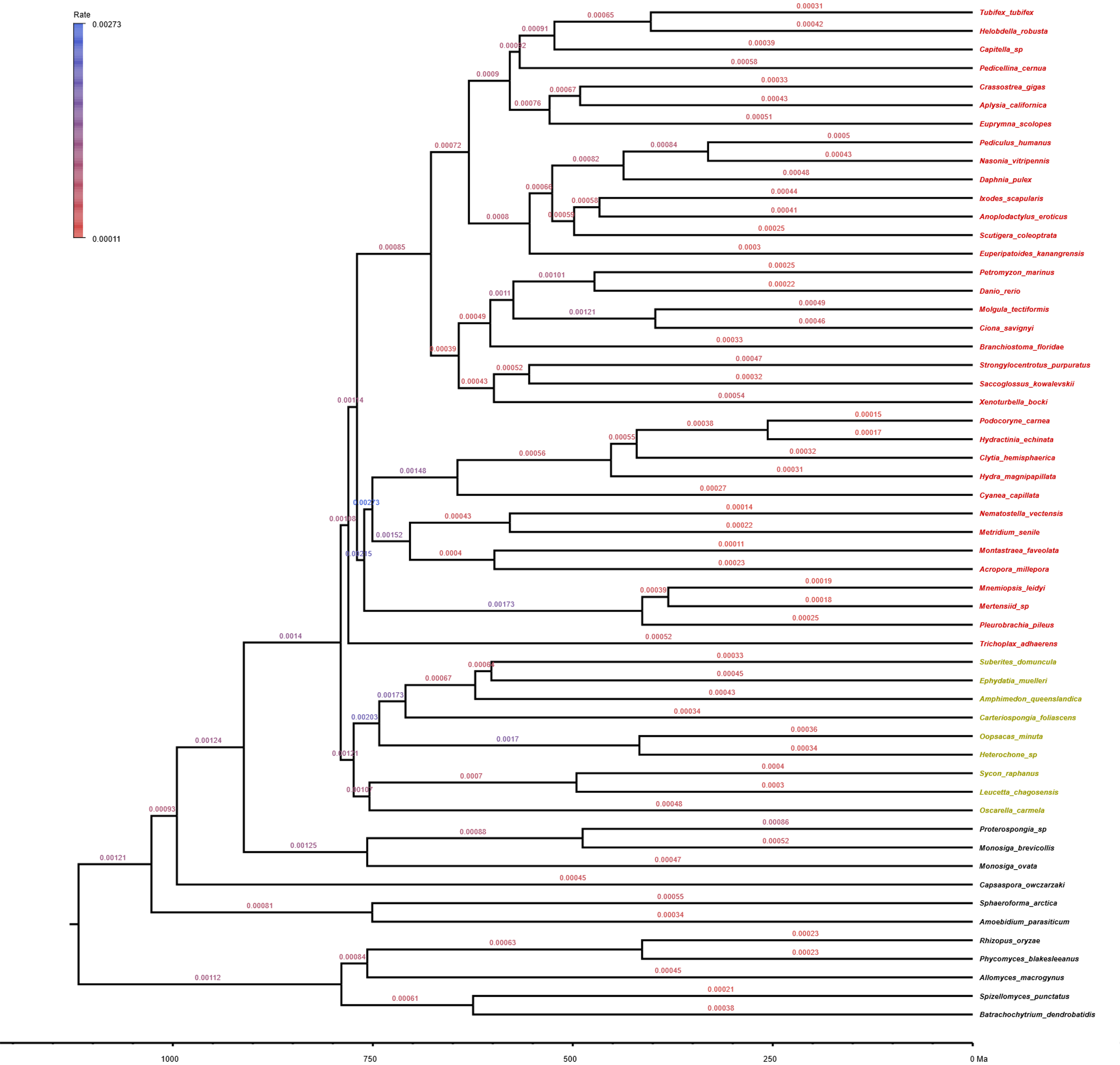

**Fig. S20.** Full chronogram showing inferred substitution rates in an autocorrelated (log-normal) relaxed molecular clock from (2) with a birth-death tree prior and root age of 1000 Ma. Branch labels show the average substitution rate (in substitutions per site per Myr) inferred along each branch. Sponges are shown in yellow, other Metazoa in red, and non-Metazoa in black.

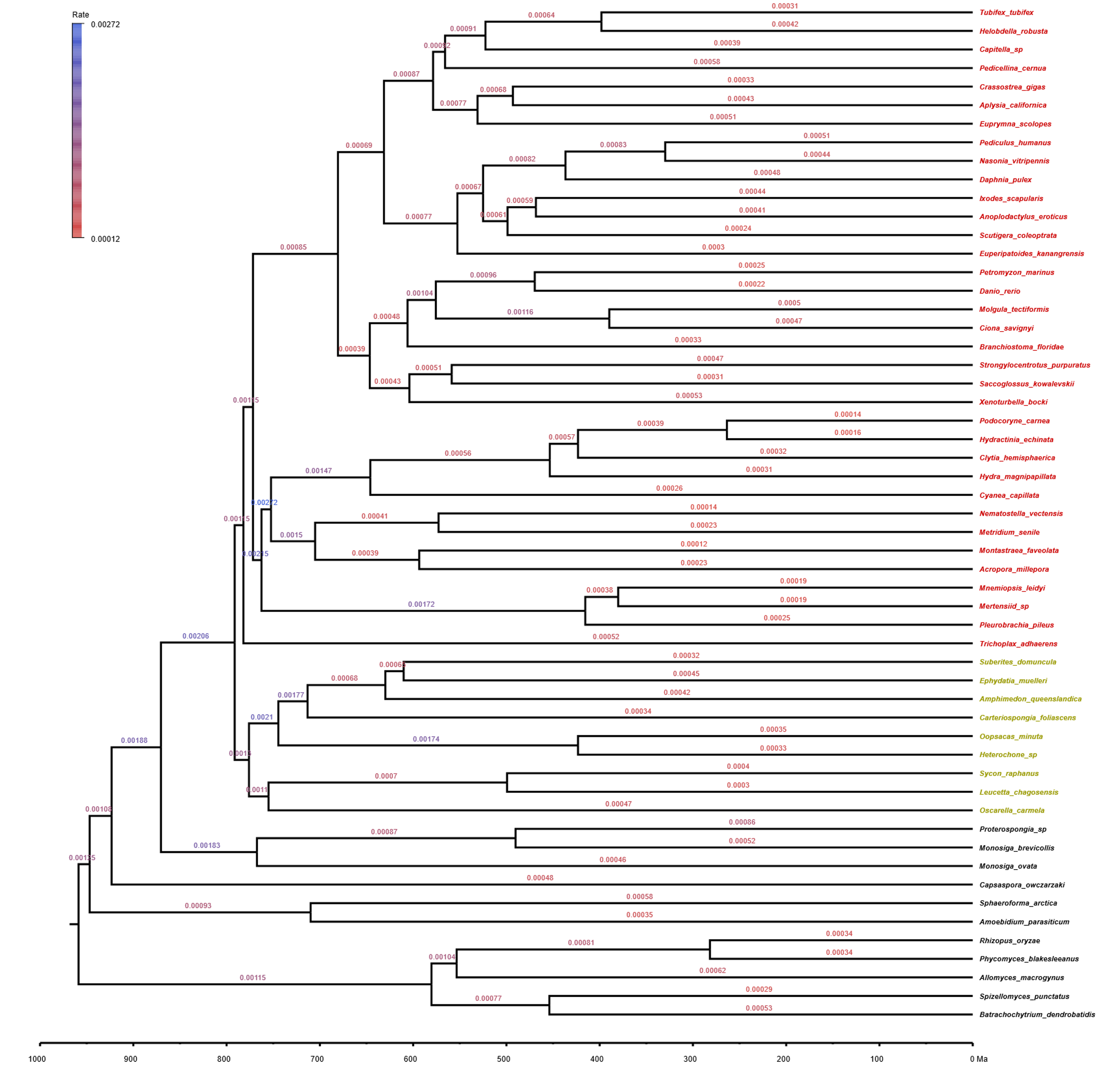

**Fig. S21.** Full chronogram showing inferred substitution rates in an autocorrelated (log-normal) relaxed molecular clock from (2) with a uniform tree prior and root age of 800 Ma. Branch labels show the average substitution rate (in substitutions per site per Myr) inferred along each branch. Sponges are shown in yellow, other Metazoa in red, and non-Metazoa in black.

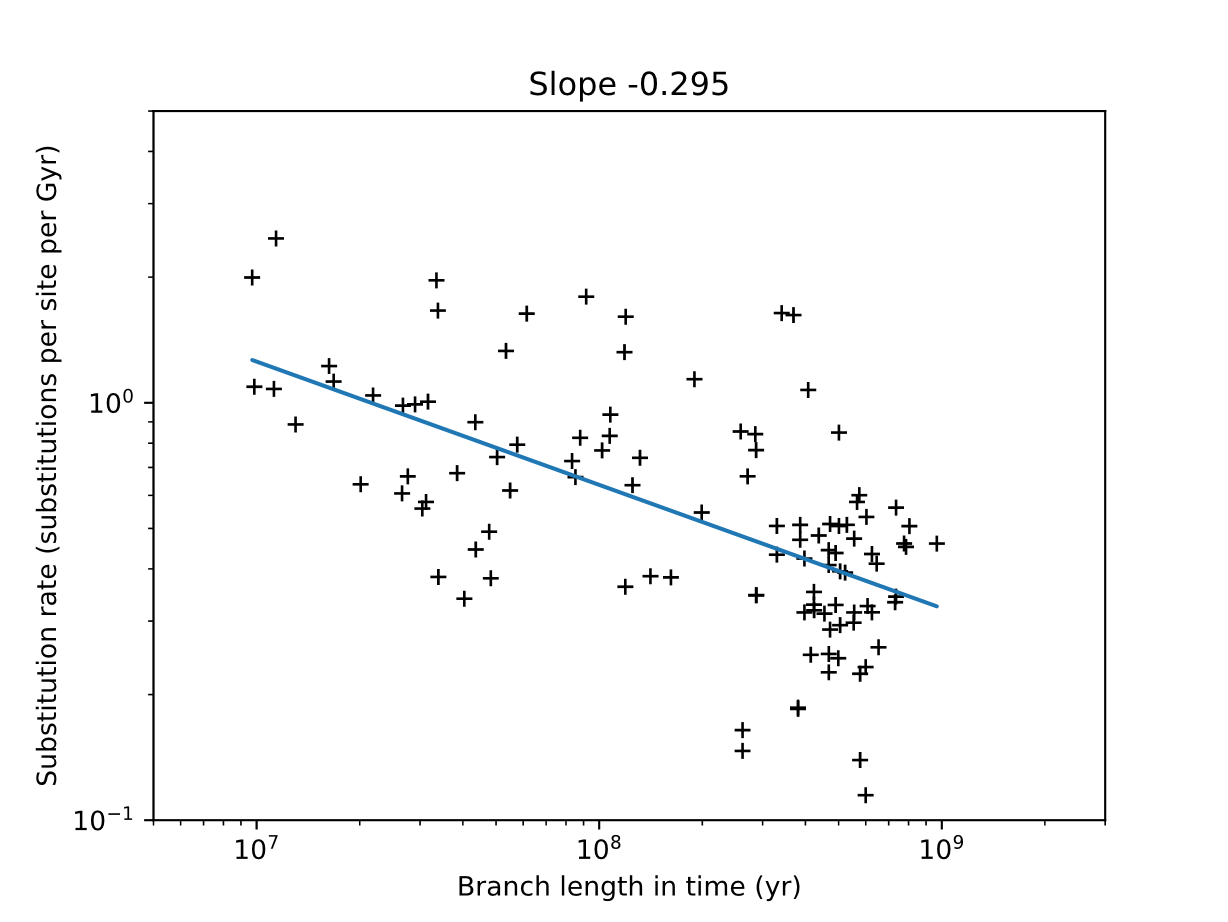

Fig. S22. The Sadler-like scale dependence of substitution rates in the dataset from (2) in a molecular clock using an autocorrelated (log-normal) clock model with a uniform tree prior and root age of 1000 Ma. The corresponding chronogram is shown in full on Figure S19.

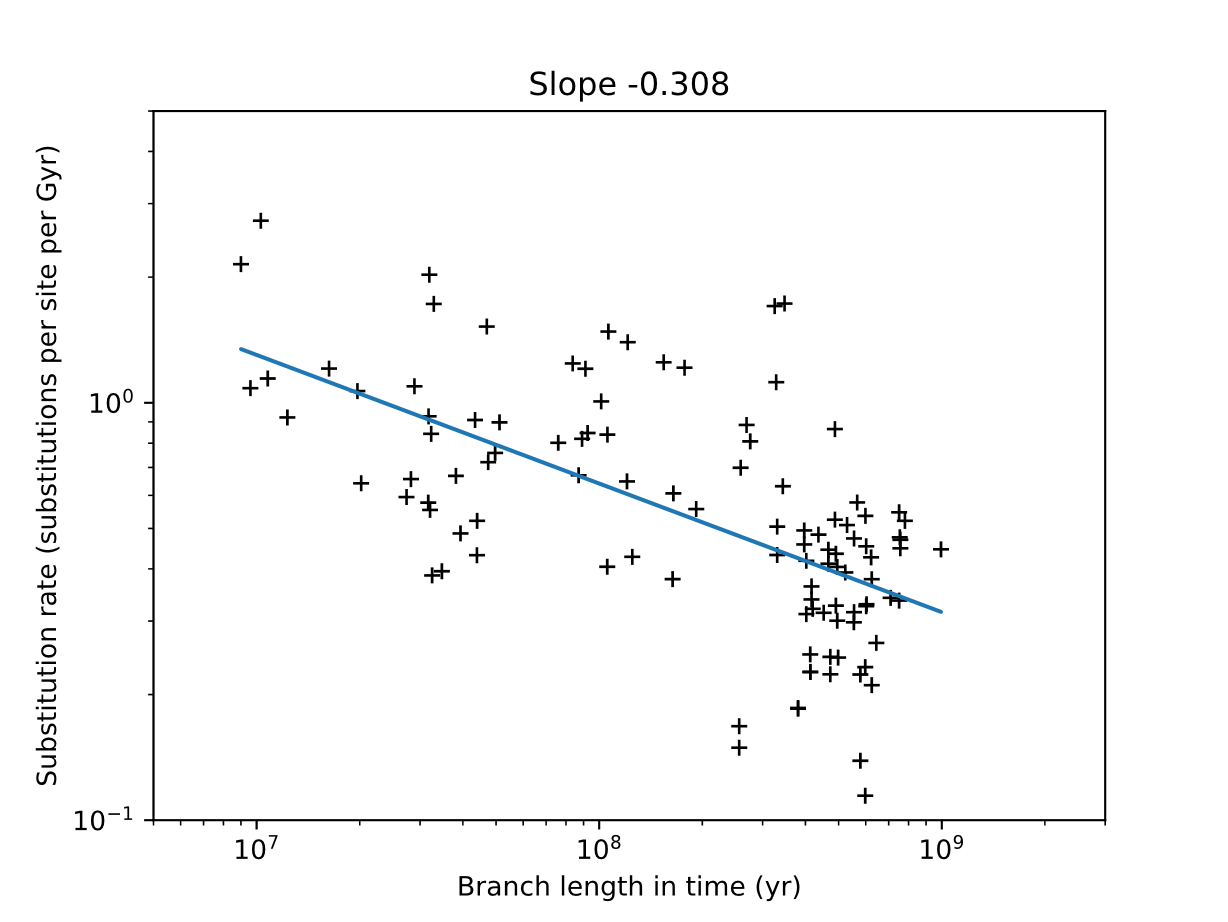

Fig. S23. The Sadler-like scale dependence of substitution rates in the dataset from (2) in a molecular clock using an autocorrelated (log-normal) clock model with a birth-death tree prior and root age of 1000 Ma. The corresponding chronogram is shown in full on Figure S20.

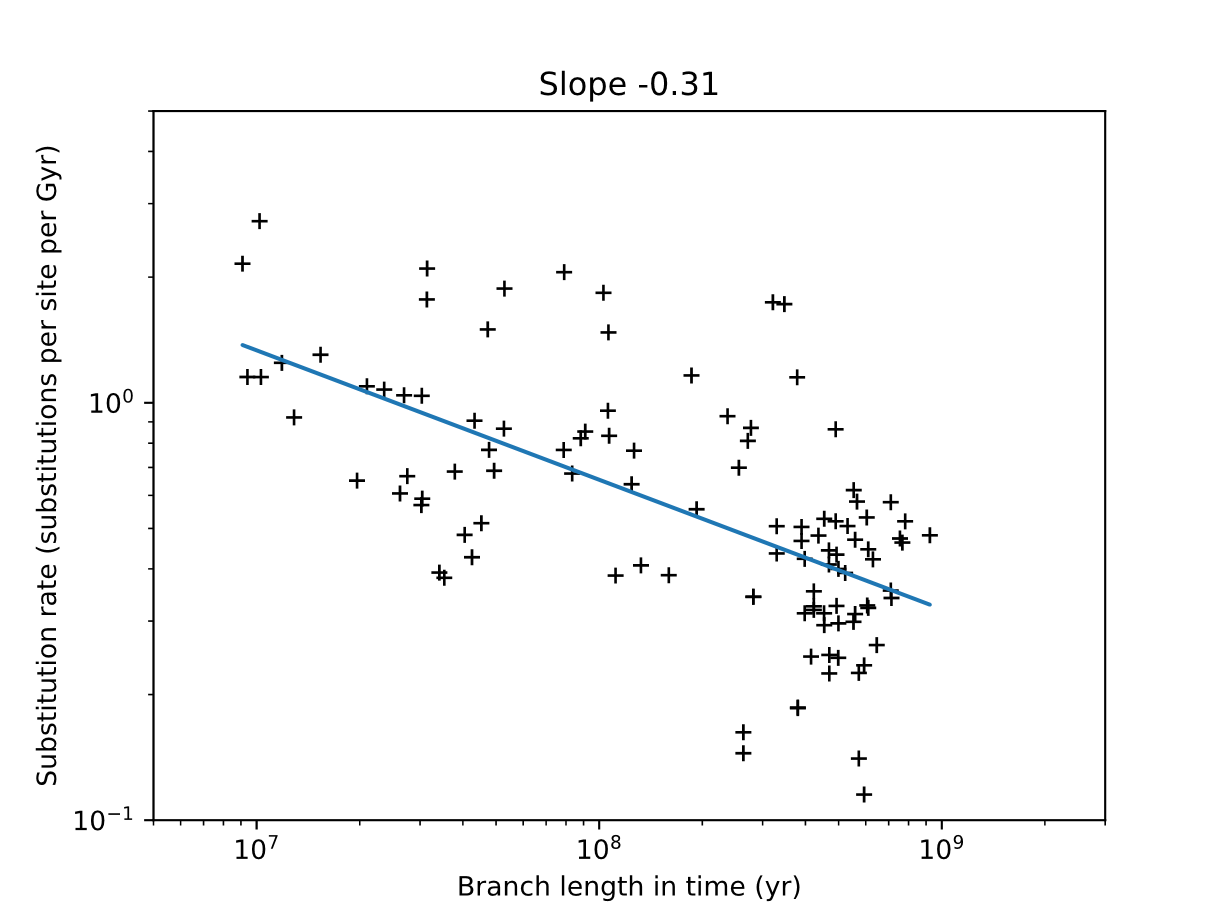

Fig. S24. The Sadler-like scale dependence of substitution rates in the dataset from (2) in a molecular clock using an autocorrelated (log-normal) clock model with a uniform tree prior and root age of 800 Ma. The corresponding chronogram is shown in full on Figure S21.
